# Sacrificial mothers: increased matrotrophy is associated with reduced maternal longevity across chondrichthyans and mammals

**DOI:** 10.64898/2026.05.07.723519

**Authors:** Rohan M Lewis, Davis Laundon

**Author notes:** Corresponding author: Davis Laundon, Faculty of Medicine, MP 887, IDS Building, University of Southampton, Southampton General Hospital, Southampton, SO16 6YD, UK.

## Abstract

Reproductive strategies vary widely among vertebrates, yet the selective drivers of life history trait evolution remain unresolved. Viviparity is typically associated with a ‘slower’ life history syndrome of larger bodies, increased lifespans, and reduced fecundity. However, viviparity is often coupled to increased matrotrophic investment, which according to the Disposable Soma Theory (DST) should reduce longevity. Conversely, the Selfish Mother Hypothesis (SMH) suggests that matrotrophic mothers withhold nutrients for the sake of their own future survival and reproduction. These opposing frameworks imply conflicting life history outcomes, and it remains unclear whether such dynamics operate primarily among individuals of the same species or shape higher-level clade-wide divergence. Here, we used a phylogenetically controlled comparative analysis of chondrichthyan (*n* = 162) and mammalian (*n* = 620) species to show that both the convergent origin and quantitative degree of matrotrophy is associated with reduced longevity across two major vertebrate clades. Our results decouple parity mode and nutrient provisioning to show that matrotrophy reduces maternal longevity, which counteracts the slower life history strategy of viviparity. We provide evolutionary support for the DST, and not the SMH, in reproductive strategy diversification. Using a simple stochastic simulation, we propose a unified ‘Sacrificial Mother’ framework, in which increased matrotrophic investment specifically reduces maternal longevity through the accumulation of somatic costs. Our work identifies the somatic costs of viviparous matrotrophy as a fundamental, but previously unrecognised, evolutionary trade-off against individual embryonic fitness which shapes the diversification of vertebrate life history strategies beyond resource allocation in intraspecific individuals.

## Introduction

There exists a remarkable diversity of reproductive strategies across the animal kingdom^1–5^. A key distinction is ‘parity mode’, which describes whether an animal reproduces by laying eggs (oviparity) or birthing live young (viviparity)^2^. The origin of viviparity is a striking example of convergent evolution, having independently arisen over 150 times in vertebrates alone from oviparous ancestors^2,6–8^. That viviparity has been so repeatedly selected for implies that it confers major fitness benefits to reproduction. Evolutionary factors hypothesised to favour the origin of vertebrate viviparity include the protection of young from predation^9,10^, expansion into previously inaccessible environmental niches^11–13^, and greater thermoregulatory and nutritive maternal control of embryonic development^14–17^. These fitness benefits represent an evolutionary trade-off against proposed costs associated with viviparous reproduction, such as constrained reproductive output, higher investment per offspring, increased predation of pregnant mothers due to burdens on locomotion, and parent-offspring conflict^18–21^.

Moving from an oviparous to a viviparous reproductive strategy necessitates changes to an animal’s life history strategy. To support internalised gestation, viviparous animals have been shown to be typically larger bodied^22–24^ and produce fewer offspring with higher individual investment per offspring^25^, although this is not always the case^26^. Higher investment offspring in viviparous species will experience individual fitness benefits from increased survival^27,28^ at the cost of fewer offspring per litter^29,30^. As death of a pregnant mother leads to offspring mortality in a way less applicable to oviparous animals with short periods of internal development, viviparous vertebrates have been shown to inhabit life history spaces or environments with lower rates of extrinsic adult mortality^31^. As larger-bodied animals, they also live longer due to the positive association of body mass and longevity^32,33^. From these studies, we can assemble a consensus ‘life history syndrome’ associated with viviparity (Figure 1). However, the oviparity-viviparity dichotomy is simplistic, and conceals a diversity of strategies to provision developing embryos with nutrients. Oviparous species provision embryos by lecithotrophy (through a finite deposited yolk source), whereas viviparous animals may be either lecithotrophic or matrotrophic (continuously provisioning embryos with maternal nutrients throughout gestation)^4^. When assessing the evolution of life history syndromes, the interaction of parity mode with nutrient provisioning, and decoupling their competing selective pressures, has rarely been considered.

**Figure 1:**
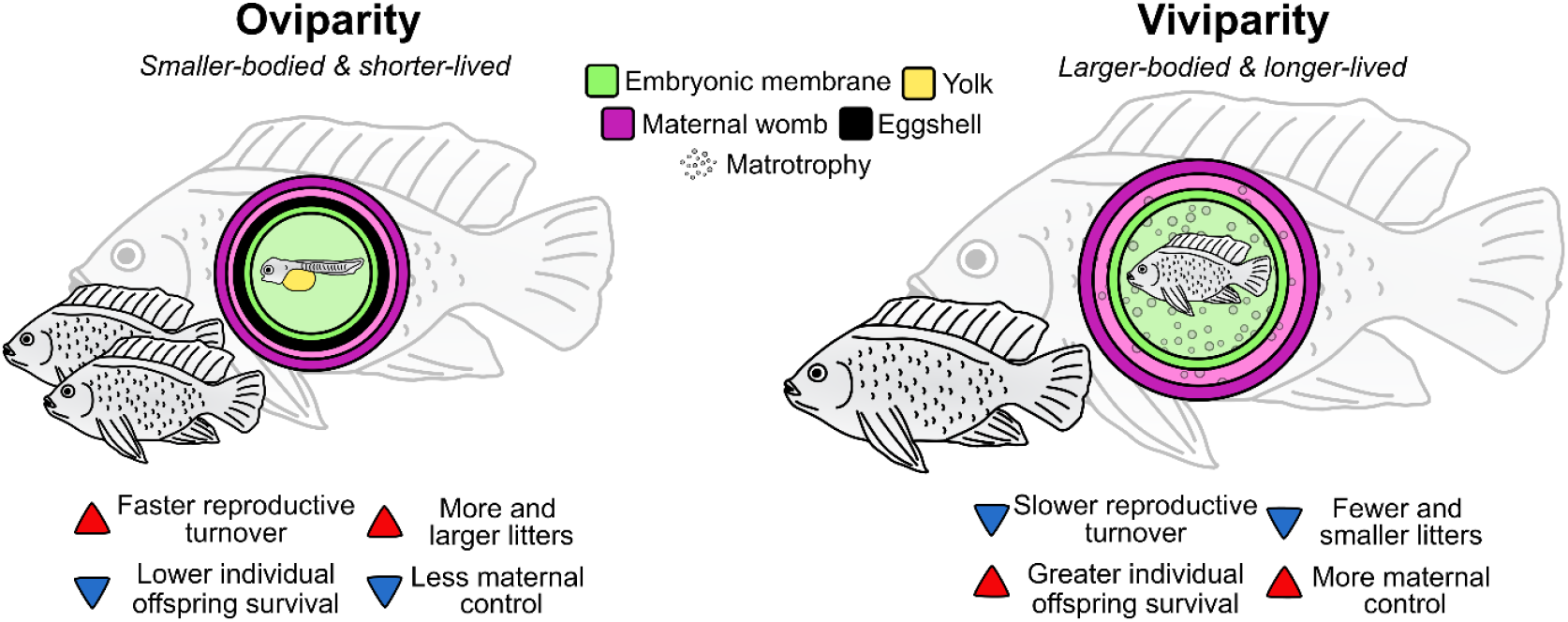
Viviparous animals are generally larger-bodied and longer-lived than their oviparous counterparts. Diagrammatic summary of the consensus understanding of the oviparous versus viviparous life history syndrome (group of associated life history traits), shown in a generalised vertebrate.

Parental animals must trade-off investment in reproduction with their own somatic maintenance. Increased investment in reproduction has been shown to reduce innate immunity^34,35^, somatic growth^36,37^, and lifespan^38,39^ within species. According to the Disposable Soma Theory (DST)^40^, aging and natural mortality of an individual are a consequence of reallocation of resources from somatic maintenance to reproduction. The DST has received mixed empirical support^41–44^. Likewise, the Trexler-DeAngelis model proposes the evolution of matrotrophy as a costly strategy which shortens maternal lifespans^45,46^. This raises an apparent contradiction in our understanding of reproductive evolution, whereby a transition to viviparity and increased reproductive investment can lead to both a larger-bodied, longer-lived life history syndrome and reduced longevity at the same time, depending on the examined variable. Indeed, it is not clear if the DST represents a macroevolutionary dynamic capable of clade-wide diversification in reproductive strategy, as investigations have mostly been intraspecific experimental studies between individuals^41,44^. Resolving how these conflicting life history variables interact is necessary to truly understand the fundamental evolutionary drivers of vertebrate reproduction.

Alternately, under the Selfish Mother Hypothesis (SMH)^17^, the greater maternal control of embryonic development conferred by viviparous matrotrophy allows a mother to withhold nutrients during hard times for the sake of her own somatic investment^47^, in a phenomenon termed ‘maternal constraint’^48^. Again, it is not clear to what extent the SMH operates beyond the intraspecific individual allocation of resources as a plastic response to changing environments. The SMH and maternal constraint are underpinned by parent-offspring conflict dynamics which state that, as the embryo is half paternal, its fitness gains for a given pregnancy can become misaligned with the long term fitness interests of the mother, leading to a tug-of-war for limited resources^19,49,50^. Parent-offspring conflict is likely a key driver of reproductive strategies such as viviparity and placentation^18,19,51–53^. Key reproductive dynamics like the DST and SMH need not be mutually exclusive, but clearly both cannot be equally dominant as evolutionary drivers as one mitigates against the other. When DST and SMH investment trade-offs are coupled to the reported viviparous life history syndrome of larger-bodied, longer-lived animals with fewer offspring, then it is unlikely that all the variables proposed to drive the evolution of viviparous matrotrophy can be simultaneously explanatory. Empirical studies into the DST and SMH typically involve comparison of animals of the same species under different experimental or environmental conditions^44,47,54^, therefore they are unable to resolve how the trade-off between investment and somatic maintenance contributes to intergroup diversification. Clade-wide investigations of these variables as evolutionary drivers remain unresolved.

Among vertebrate groups, chondrichthyans and mammals represent powerful complimentary systems to investigate the evolution of reproductive strategies and maternal investment. Chondrichthyans (sharks, skates, rays, and chimaeras) are a large and ancient (~420 Ma) clade which exhibits the widest range of reproductive strategies among vertebrates, where viviparity has evolved independently at least 12 times and matrotrophy has evolved at least 6 times, with all known matrotrophic strategies being present^4,7,55^. The wide diversity and repeated convergence of chondrichthyan reproduction makes the group a compelling natural system to investigate the evolutionary origins of viviparous reproductive strategies. Conversely, mammals are unique amongst vertebrates in that every species (except monotremes) are viviparous and matrotrophic, with this strategy having originated only once in their shared common ancestor ~130 Ma^7,56–58^. Yet, within this single strategy there is a wide divergence in the levels of maternal investment between species from the ancestral condition^59,60^. Total maternal investment can now be quantified between mammal species, allowing for recent systematic evolutionary investigations into the variation of viviparous matrotrophy across the whole group^53,61^. A combined investigation of both the chondrichthyans and mammals would be a powerful natural experiment to uncover changes to key life history traits associated with both the origin and degree of viviparous matrotrophy, under convergent and divergent evolutionary scenarios, such as the unclear relationship between adult longevity and reproductive investment.

Here, we investigated the association between reproductive strategies and life history traits in chondrichthyans and mammals, with a focus on the relationship between embryonic investment and maternal longevity. For both groups we compiled large cross-species datasets of adult body size, lifespan, and reproductive data. For chondrichthyans, we used information on parity mode and nutrient provisioning^4^ to compare the independent origin of different reproductive strategies across species, and for mammals we used a quantitative metric^61^ to examine different levels of maternal investment within an exclusively viviparous matrotrophic clade with a single evolutionary origin. For both chondrichthyans and mammals, we built a phylogenetic tree for all the species in our compiled datasets from well-established phylogenies^62,63^. As such, we could control for both the effect of overall body size on longevity and phylogenetic distance between species, to robustly investigate the interaction of maternal investment and longevity. We hypothesised that, when controlled for absolute body size and phylogeny, both the adoption of viviparous matrotrophy in chondrichthyans and the degree of investment in mammals would be associated with reduced longevity due to the trade-offs between reproduction and somatic maintenance across species. Using a simple stochastic simulation, we go on to propose an evolutionary scenario of how viviparous matrotrophy can lead to a reduction in longevity at the scale of clade-wide divergence in life history strategies.

## Methods

### Dataset collation and phylogenetic tree construction

Life history metrics for chondrichthyans were compiled using previously published datasets from^64–66^ (*Supplementary File 1*). The aggregated data was filtered for maximum adult body length (cm) and maximum longevity (years) (using female-specific data where available), which were the best covered size and longevity metrics. Data were manually checked and updated where more recent values were available using Sharkipedia^67^, FishBase^68^, and the IUCN Red List of Threatened Species. Data for chondrichthyan parity modes and nutrient provisioning strategies were then added to the dataset from^4^. For placental mammals, adult mass (kg) and lifespan (days) were taken from the dataset compiled by^61^ used in^53^ (*Supplementary File 2*). The maternal investment from a multiple phylogenetic linear mixed model (*MI*_*MPLMM*_) metric created by^61^ to quantify maternal investment in mammals was taken from the same dataset. Using the species for which life history data was present, phylogenetic trees were built from the online resource https://vertlife.org/phylosubsets/ based on published chondrichthyan^62^ and mammalian^63^ phylogenies. A consensus tree for each group was generated from 1,000 tree replicates using *consensus*.*edges* in *phytools*^69^ run with R v.4.4.0 implemented in RStudio v. 2025.09.2. Genus names for species with life history data were updated to match the tip labels of the consensus tree where differently assigned in the phylogeny.

### Data analysis

Body size influences lifespan beyond reproductive strategy^32,33^, which may confound analysis. Likewise, the species in our datasets are not independent data points but related by phylogenetic proximity, which may also explain variations in life history beyond reproductive strategy. Therefore, the influence of body size and phylogeny on longevity was investigated and controlled for. For both datasets, body size and longevity data were log transformed and initially fitted with a simple non-phylogenetic linear regression model (LM) to account for the allometric scaling of lifespan with adult size (*Supplementary File 3*). Then, a phylogenetic generalized least squares (PGLS) model was implemented using the *gls* function in *nlme* with a Pagel’s λ correlation structure to factor in phylogenetic distance between species. PGLS models were initially fitted using maximum likelihood (ML) for comparison purposes and then final parameters were derived with restricted maximum likelihood (REML) models. The fits of phylogenetically corrected PGLS models to the data were compared to non-phylogenetic linear regression models using Akaike Information Criterion (AIC) scores. Residuals from the allometric scaling of longevity by body size using a PGLS model were extracted for downstream analysis, and their distributions were visually inspected using histograms.

Chondrichthyan species were categorised into three reproductive strategies based on parity mode and nutrient provisioning as described by^4^: oviparous lecithotrophy, viviparous lecithotrophy, and viviparous matrotrophy. Longevity residuals between reproductive strategies were compared using a Kruskal-Wallis test with Dunn’s post hoc to explore if reduced longevity beyond the influence of body size and phylogeny was associated with different reproductive strategies. For mammals, *MI*_*MPLMM*_ (a continuous variable to quantify reproductive investment) was analysed for relationship against size-weighted longevity using non-phylogenetic (Pearson’s correlation) and PGLS models as above and compared for the best fit. For both groups, PGLS longevity residuals were mapped onto the consensus phylogenetic tree using the *phytools* function *contMap*, and an outer ring representing either reproductive strategy (chondrichthyans) or *MI*_*MPLMM*_ values (mammals) was overlaid, visualising the phylogenetic spread of maternal investment with relative longevity. All analysis was conducted using the packages *tidyverse, ape, nlme*, and *phytools* run with R v.4.4.0 implemented in RStudio v. 2025.09.2.

### Simple stochastic simulation of maternal damage

We developed a simple stochastic simulation of biological variables that we propose increase maternal damage in matrotrophic species and contribute to reduced longevity as a conceptual evolutionary scenario. We modelled three reproductive strategies: oviparous lecithotrophy, viviparous lecithotrophy, and viviparous matrotrophy. Simulations were run for 100 generations from an initial level of embryo investment for replicate populations (*n* = 100 per strategy) with a fixed random seed to allow reproducibility. Embryonic investment encompasses both provisioning and developmental support without necessarily implying a direct increase in physical dimensions. Embryonic investment evolved through iterative mutation and selection interaction. For lecithotrophic strategies, investment was constrained by an upper ceiling representing the limits of resource allocation from a finite deposited yolk source. Viviparous lecithotrophy was bound by this ceiling, however it was permitted a small increase to accommodate the increased investments of internalised gestation, such as maternal thermoregulation. Matrotrophy exceeds the yolk threshold by investing additional maternal resources.

Embryo fitness was modelled as a function of embryonic investment, with diminishing returns on fitness at higher levels of investment and a small increase for viviparous strategies to reflect maternal protection and developmental control of internalised embryos. Matrotrophy accrued additional fitness by investing nutrients beyond the yolk threshold. At each generation, embryos with higher fitness were preferentially retained, while less-fit embryos could persist at a probabilistic rate, allowing stochastic exploration of the trait space. Maternal damage was modelled as a non-linear function of excess investment above a defined damage threshold, reflecting somatic costs on the mother from high maternal investment, such as resource drain or biomechanical damage from obstetric problems. Once embryo investment exceeded the damage threshold, maternal damage increased according to *maternal damage = s × (e)*^*d*^ where *s* = the scaling component, *e* = the degree investment exceeds the threshold, and *d* = the damage exponent. Lecithotrophic species did not incur maternal damage as their investment did not exceed the damage threshold. Maternal lifespan was reduced as a non-linear function of accrued maternal damage, with large longevity costs for more severe damage. Baseline maternal lifespan differed between reproductive strategies, with viviparous strategies assigned higher initial lifespans to reflect their association with larger-bodied, lower extrinsic mortality life history syndromes. However, only matrotrophic strategies experienced reductions in lifespan because of increased maternal damage. All simulation was conducted in Python 3.12.7 implemented in JupyterLab 4.2.5. Portions of code used in this study were generated with the assistance of ChatGPT-5.2 model (OpenAI), then reviewed and curated by the authors. The full stochastic simulation code with adjustable parameters is available as *Supplementary File 4*.

## Results

We compiled datasets containing adult body size, lifespan, and reproductive strategy or maternal investment for chondrichthyan (*n* = 162) (Figure 2&3) and mammalian (*n* = 620) (Figure 4&5) species. In absolute terms, oviparous lecithotrophic chondrichthyans were smaller than both viviparous lecithotrophic (*p* < 0.05) and viviparous matrotrophic species (*p* < 0.01), and viviparous lecithotrophic chondrichthyans were longer lived than both oviparous lecithotrophic (*p* < 0.05) and viviparous matrotrophic species (*p* < 0.05) (Figure 3A&B). In mammals, there was no significant relationship between absolute body size and maternal investment (*p* = 0.66), however mammals which invested more in reproduction had shorter lifespans (*p* < 0.01) (Figure 5A&B). Lifespan was positively correlated with body size in both chondrichthyans (LM R^2^ = 0.35, *p* < 0.001) (Figure 3C) and mammals (LM R^2^ = 0.49, *p* < 0.001) (Figure 5C), and therefore longevity was corrected by body size prior to comparison with reproduction. A phylogenetic PGLS model was found to fit the data better than a simple linear regression for both chondrichthyans (ΔAIC 59.1, Pagel’s λ = 0.82) and mammals (ΔAIC 655.2, Pagel’s λ = 0.99) and was therefore used to calculate longevity residuals representing deviations in life span relative to that predicted from body size and controlled for phylogenetic distance.

**Figure 2:**
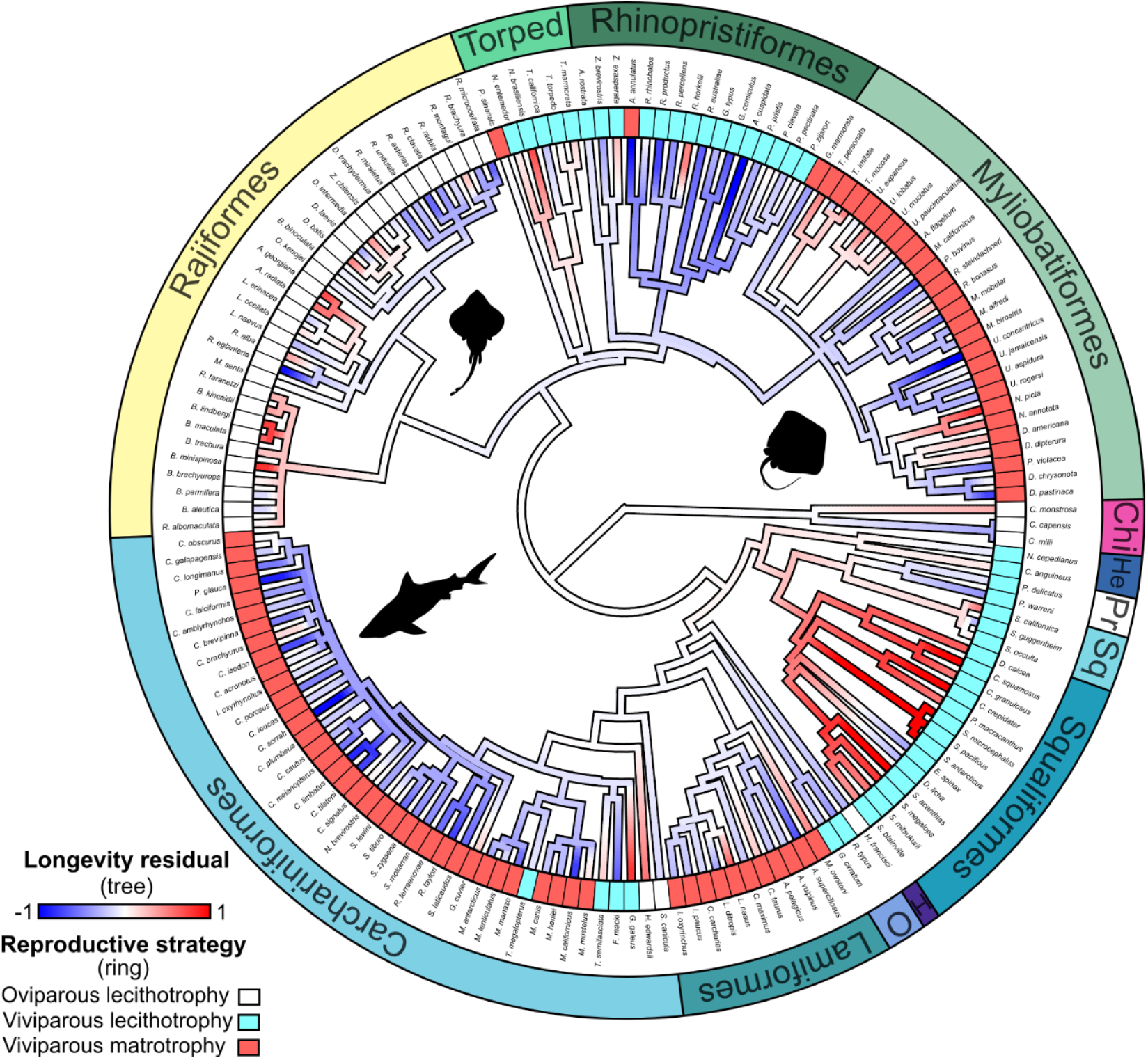
Clades of chondrichthyans which have evolved viviparous matrotrophy exhibit lower relative longevity. Phylogenetic distribution of chondrichthyans used in this study (*n* = 162), showing the relative longevity (tree) and reproductive strategy (ring) for each species. Orders labelled on the outer most ring. Chi = Chimaeriformes, H = Heterodontiformes, He = Hexanchiformes, O = Orectolobiformes, Pr = Pristiophoriformes, Sq = Squatiniformes, Torped = Torpediniformes. Silhouettes from https://www.phylopic.org/ signpost major groups.

**Figure 3:**
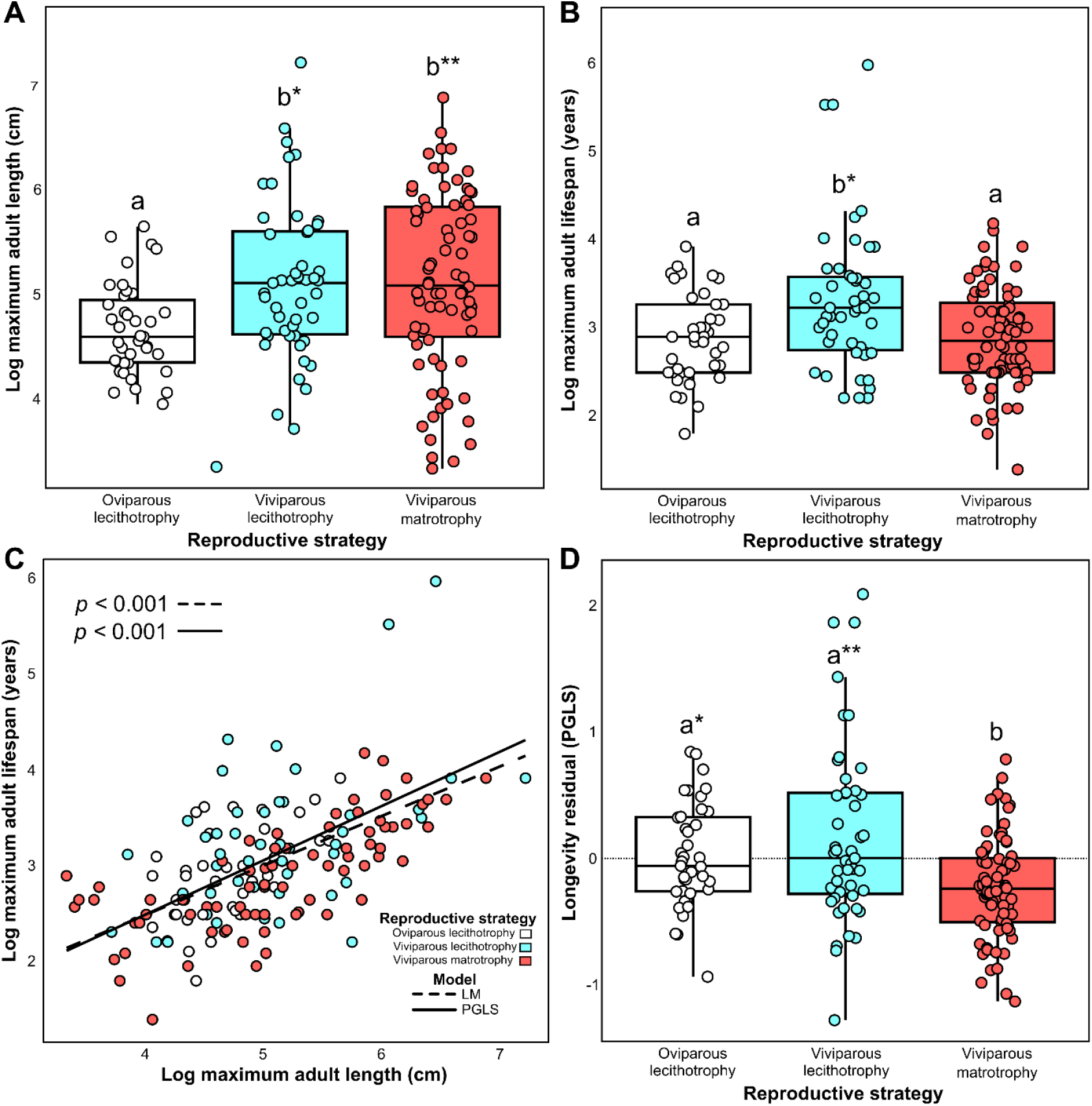
Viviparous matrotrophy is associated with reduced longevity in chondrichthyans. Viviparous species were larger than oviparous species (A) and viviparous lecithotrophic species lived longer than either matrotrophic or oviparous species (B). (C) Adult body size was positively correlated with lifespan in chondrichthyans, and a phylogenetically corrected model (PGLS) fit the data better than a simple regression (LM) (ΔAIC 59.1, Pagel’s λ = 0.82). (D) Comparison of size-corrected and phylogenetically controlled longevity (longevity residual) between different chondrichthyan reproductive strategies. Significant differences between reproductive strategies are denoted by different letters. \**p* < 0.05, \*\**p* < 0.01.

**Figure 4:**
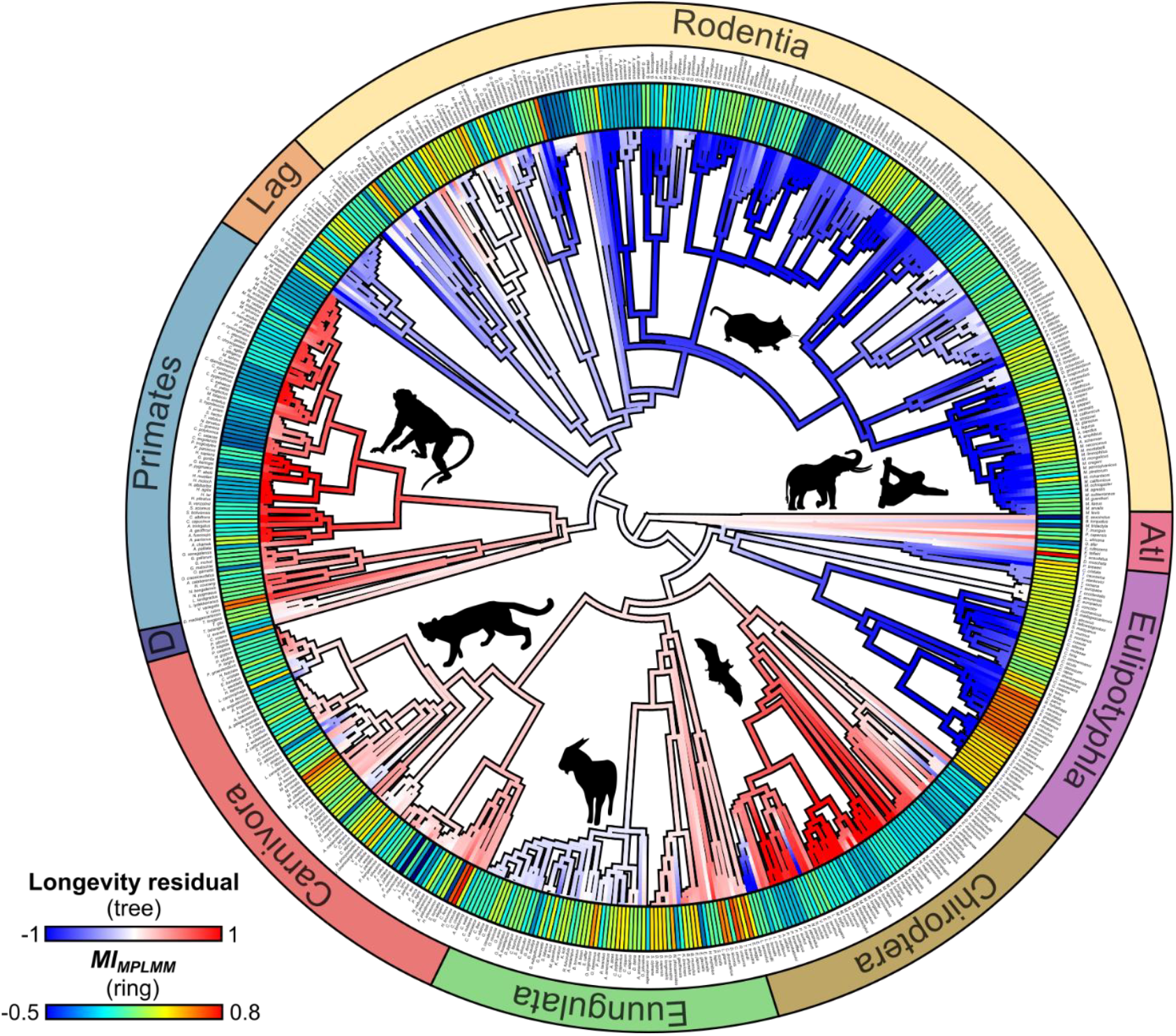
Mammalian clades with high maternal investment are shorter-lived. Phylogenetic distribution of placental mammal species used in this study (*n* = 620), showing relative longevity (tree) and maternal investment (*MI*_*MPLMM*_, ring). Orders labelled on the outer most ring. Atl = Atlantogenata, D = Dermoptera, Lag = Lagomorpha. Silhouettes from https://www.phylopic.org/ signpost major groups.

**Figure 5:**
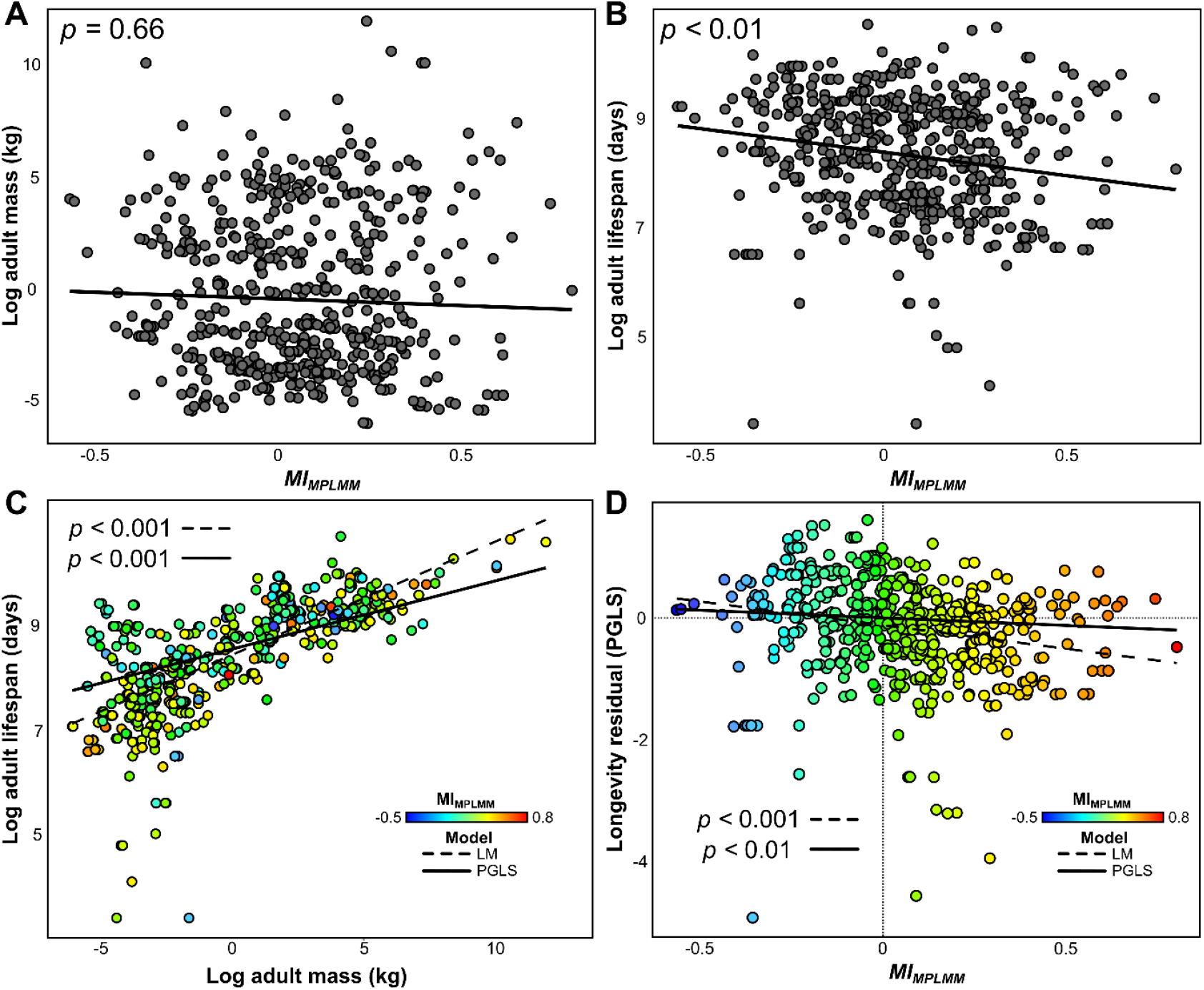
Mammal species that invest more in reproduction live shorter lives. Maternal investment (*MI*_*MPLMM*_) showed no association with body size (A) but was negatively associated with lifespan (B). (C) Adult body size was positively correlated with lifespan in mammals, and a phylogenetically corrected model (PGLS) fit the data better than a simple regression (LM) (ΔAIC 655.2, Pagel’s λ = 0.99). (D) Maternal investment was negatively associated with size and phylogeny corrected longevity.

Overlaying longevity residuals onto the consensus trees lets us inspect the phylogenetic distributions of relative lifespan and reproductive investment across clades (Figures 2&4). In chondrichthyans (Figure 2), majority viviparous matrotrophic orders have shorter lifespans than would be predicted by allometry (e.g Carcharhiniformes (Longevity residual) = −0.30 ± 0.06 mean ± SEM; Myliobatiformes; −0.10 ± 0.08; Lamniformes = −0.09 ± 0.11) whereas majority viviparous lecithotrophic orders are relatively longer lived (such as the superorder Squalomorphi = +0.50 ± 0.06; Torpediniformes = +0.11 ± 0.13), with oviparous lecithotrophic groups lying in between (e.g Rajiformes +0.03 ± 0.07). The notable exception is the Rhinopristiformes, a majority viviparous lecithotrophic order which has a shorter longevity than predicted by allometry (−0.39 ± 0.11). In mammals (Figure 4), orders associated with higher maternal investment^53,61^ tend to have lower size-corrected longevity (Eulipotyphla MI = +0.36 ± 0.02, Longevity residual = −0.88 ± 0.08; Lagomorpha MI = +0.07 ± 0.05, Longevity residual = −0.384 ± 0.074; Rodentia MI = +0.04 ± 0.01, Longevity residual = −0.70 ± 0.05). Inversely, orders which underinvest in reproduction live longer than would be expected (Primates MI = −0.11 ± 0.02, Longevity residual = +0.73 ± 0.03; Chiroptera MI = −0.12 ± 0.01, Longevity residual = +0.65 ± 0.09) with other groups in between. Overall, viviparous matrotrophic chondrichthyans had a lower relative longevity than both viviparous lecithotrophic (*p* = 0.02) and oviparous lecithotrophic species (*p* = 0.002) (Figure 3D). In mammals, maternal investment (*MI*_*MPLMM*_) was inversely proportional to relative longevity (PGLS *p* = 0.005) (Figure 5D) indicative of a direct relationship.

Using a simple stochastic simulation, we illustrate a scenario of how biological variables associated with viviparous matrotrophy could interact to reduce maternal longevity between life history strategies (Figure 6). We propose that maternal damage, which reduces maternal longevity, is a non-linear consequence of high maternal investment associated with viviparous matrotrophy. The degree of maternal investment per embryo is constrained in lecithotrophic strategies by the limit of resources from a deposited yolk without supplemental matrotrophic investment (Figure 6A&B), and as such maternal damage is mitigated. We give viviparous lecithotrophy a small fitness increase due to the protection and control of internalised embryos, but we propose it is still fundamentally constrained by the finite limit of deposited yolk resources. In contrast, viviparous matrotrophy can exceed this ceiling by provisioning with additional maternal nutrients, allowing embryos to achieve higher levels of investment and individual fitness (Figure 6B). We propose that once embryo investment passes a damage threshold, whereby high embryonic investment is detrimental to the mother, maternal damage increases non-linearly (Figure 6C). This elevated investment confers high fitness increases to the embryo but reduces maternal longevity as a function of accumulated costs (Figure 6D). High matrotrophic investment therefore represents a trade-off with the increased fitness of individual embryos conferred by higher investment strategies against maternal somatic maintenance. We summarise the nature of these distinct costs of viviparous matrotrophy, where maternal longevity is a direct trade-off against reproductive investment, underpinning our ‘Sacrificial Mother’ model of reproductive strategy evolution (Figure 6E).

**Figure 6:**
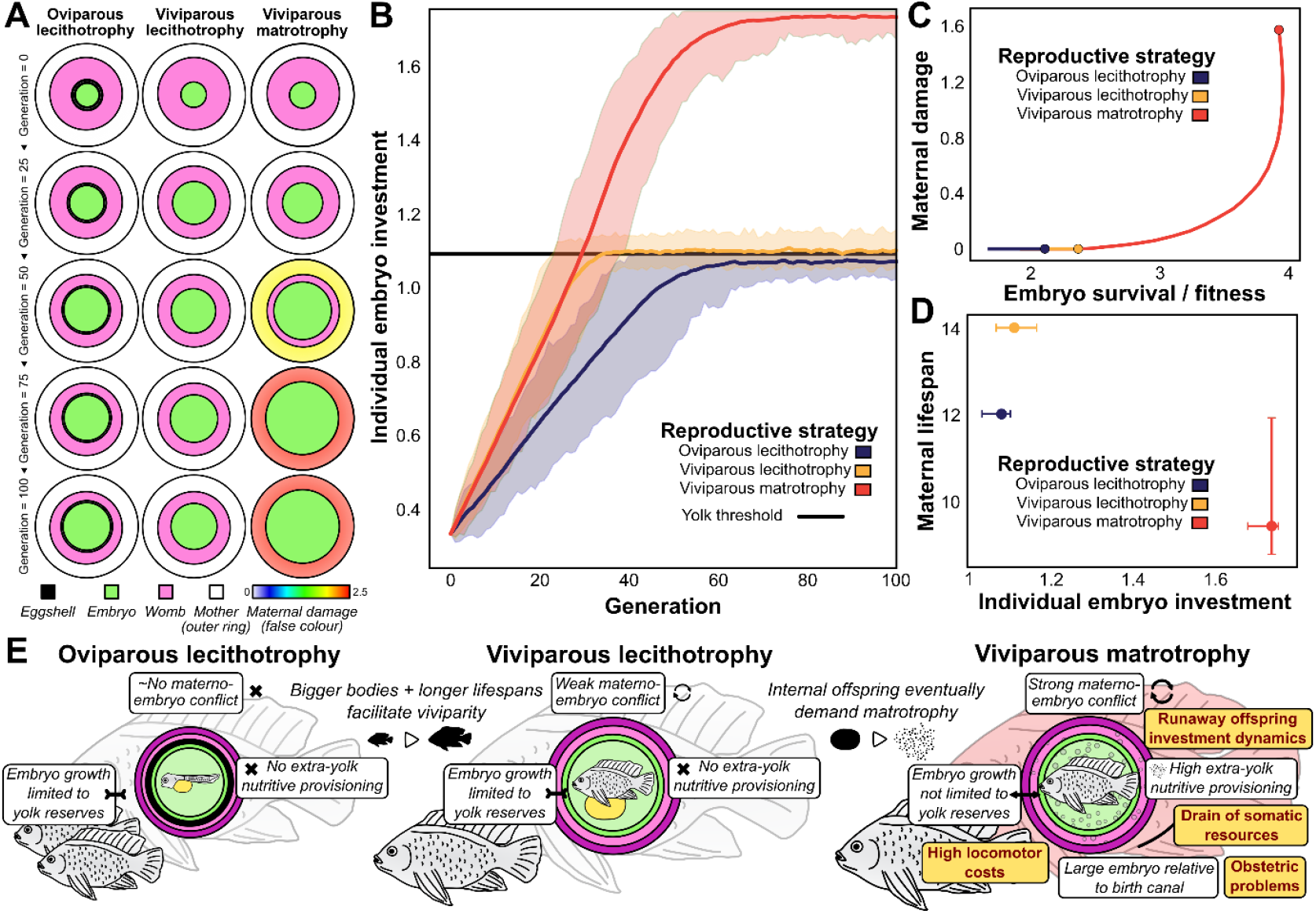
We propose a ‘Sacrificial Mother’ model for vertebrate reproductive evolution where maternal longevity is reduced by non-linear costs of matrotrophy. (A) Visual representation of a simple stochastic simulation of embryonic investment and maternal damage for different reproductive strategies on a single lineage trajectory. The size of the embryo reflects the degree of investment, not necessarily the physical size of the offspring. We use ‘womb’ as a general term for the maternal reproductive chamber in any species. The womb size remains fixed as a visual representation of a maternal investment threshold, which when approached or exceeded in viviparous matrotrophy, induces the onset of non-linear maternal damage (false coloured). (B) Simulated embryo investment over 100 generations for different reproductive modes. We propose the nutrients in a finite yolk desposit as the upper ceilings to embryo investment for both oviparous and viviparous lecithotrophy, with viviparity conferring a small boost from embryo protection. (C-D) Our model predicts maternal damage as a non-linear consequence of increased embryonic investment, which trades off higher individual embryonic fitness (C) against reduced maternal longevity (D). All units in A-D are arbitrary. (E) Diagrammatic summary of key costs of viviparous matrotrophy which reduce maternal longevity under our ‘Sacrificial Mother’ model, shown in a generalised vertebrate.

## Discussion

Here, we compiled large datasets of life history metrics and reproductive strategies for two diverse vertebrate clades to show that increased maternal investment is associated with reduced maternal longevity at the macroevolutionary scale. In chondrichthyans, viviparous species were bigger than oviparous ones, regardless of nutritive provisioning mode, in line with the proposal that viviparity is associated with larger body sizes^22–24^. However, only viviparous lecithotrophic species exhibited longer lifespans, with matrotrophic species living similar lengths as oviparous ones, suggesting that the positive association of larger body sizes and longer lifespans with viviparity^32,33^ is countered by the negative effect of reproductive investment on longevity^39,40^. We analysed size-corrected and phylogenetically controlled longevity residuals and found that viviparous matrotrophic chondrichthyans live shorter lives than would be predicted by allometry. Our analysis supports a DST interpretation of chondrichthyan reproductive evolution at a cross-species level, whereby mothers trade off increased matrotrophy with reduced somatic maintenance. In mammals, we found that body size did not change with the degree of maternal investment, which is in keeping with the proposed relationship between parity mode and body size as all tested mammals were viviparous. However, overall lifespan decreased with increased maternal investment, and relative longevity decreased significantly with increased maternal investment even when controlled for body size and phylogeny. As in chondrichthyans, our analysis of reduced longevity as a function of increased maternal investment supports DST dynamics beyond individuals and as a driver of high-level reproductive evolution.

Our clade-wide analysis in support of DST is contrary to some empirical studies into mammal reproductive investment, which have not found support for disposable soma models at the intraspecific level^41,70^. Our analysis also does not support the SMH, at least not at the scale of the clade across the total lifespan, contrary to empirical support for selfish mother dynamics in mammals at the intraspecific level under resource-limited conditions^54,71^. While we do not disagree that maternal constraint is adaptive to individual viviparous matrotrophic mothers in resource scarce environments^48^, our analysis finds no support that maternal control can counter the longevity reducing effects of maternal investment predicted by the DST and parent-offspring conflict ^19,53^ at the level of cross-species life history strategy evolution and clade diversification. Combined, our analysis supports the DST, with no clear support for the SMH, in two large and disparate clades of vertebrates, uniquely associated with the origin and degree of viviparous matrotrophy. As such, we propose the ‘Sacrificial Mother’ explanatory framework for clade-wide diversification of life history strategies, beyond the previously proposed intraspecific application of the DST, whereby maternal longevity is reduced due to distinct costs to somatic maintenance incurred by high viviparous matrotrophy.

Using a simple stochastic simulation, we outline how these variables may interact to reduce maternal longevity in viviparous matrotrophic species. An oviparous species that transitions to viviparity, either to expand into new environmental niches or to increase maternal prepartum developmental control or protection of young^10,11,13,14^, moves towards a larger-bodied and longer-lived life history space^23,32,33^. This produces fewer offspring with higher individual investment^25,31,72^, moving viviparous lecithotrophic species K-wards along the *r*-K continuum^73,74^. So long as a viviparous species remains lecithotrophic, it can continue to occupy a slower life history space of lower extrinsic mortality and increased longevity due to the lack of high maternal damage. Under our Sacrificial Mother model, only the origin and intensification of matrotrophy in a viviparous clade counteracts maternal longevity, due to the trade-off between its uniquely intense reproductive investment and somatic maintenance, which offsets the larger-bodied longer-lived life history space of viviparity by itself. As such, Sacrificial Mothers exhibit a distinct life history syndrome composed of both the production of highly *K-*selected offspring and a faster maternal life history strategy, relative to their lecithotrophic cousins.

The uniquely intense trade-off between reproductive investment and somatic maintenance in viviparous matrotrophic species is just one variable that we propose contributes towards reduced maternal longevity. Maternal investment in viviparous matrotrophic animals is high, often including post-partum lactation and parental care beyond direct gestational investment, which can all contribute to decreased maternal longevity^75–77^. Additionally, mothers can experience direct reductions in individual fitness due to the locomotory burdens of pregnancy^78,79^ and from morbidity and mortality from obstetric complications^41,80^. Such costs are fundamentally a result of the heightened parent-offspring conflict in viviparous animals^19,51,81^. As sexually reproduced offspring are half paternal, their fitness interests can be misaligned with that of the mother for any given pregnancy^51,53,82^, which is particularly pronounced for mostly non-monogamous clades like chondrichthyans and mammals^83,84^. Due to paternal influence, it may be in the fitness interest of a given embryo to extract more nutrients from the mother than are optimal for her to give considering her long-term fitness incorporating future reproductive events, and as such paternal genes have been shown to have a disproportional effect on increased embryonic growth during pregnancy^51,82,85^.

Mothers may counter this effect through maternal constraint in limited circumstances^48^, but our analysis shows this does not counter excessive embryonic demand as an evolutionary driver of reduced longevity across large groups. Bidirectional tissue-tissue interactions between embryo and mother are limited in oviparous species by the physical presence of an eggshell^86,87^ and short length of internalised development, and so the mechanisms of parent-offspring conflict are hindered. In viviparous species, the absence of an eggshell permits novel innovations of tissue-tissue interactions, however parent-offspring conflict will be weak in the absence of extra-yolk maternal nutrients. We propose here that lecithotrophic viviparity evolves primarily because of increased maternal protection (e.g. from predation) and developmental control of embryos as part of a slower, K-shifted life history strategy, however the novel intimacy of materno-fetal tissues and longer internalised gestation generates a runaway conflict dynamic where embryos manipulate maternal physiology to extract additional nutrients^53,88^. As such, we hypothesise that matrotrophy initially emerges and strengthens because of embryonic demand, and not maternal supply.

Our study is the first large scale analysis of body size, longevity, and reproductive strategy in chondrichthyans and mammals, using the best data currently available. Our findings will be constrained by the amount and quality of data available for sampled taxa, especially for chondrichthyans. It is likely that maximum recorded length and lifespan, although the best and most numerous data presently available, are less robust proxy metrics for size and longevity in chondrichthyans than mean adult mass and mean lifespan sampled from many individuals. In addition, we can currently only categorise matrotrophic investment qualitatively in chondrichthyans, which is less informative than standardised quantitative metrics such as *MI*_*MPLMM*_. Due to their marine habitat, relative rarity, and husbandry difficulties, life history data for chondrichthyans will likely always lag behind that for mammals. However, using the best data currently available our results demonstrate a significant reduction in maternal longevity associated with the adoption of viviparous matrotrophy in chondrichthyans and we suggest this finding will only strengthen as more and better life history metrics become available. Likewise, for both chondrichthyans and mammals, adult body size and longevity are much more readily available than female-specific data, which is more useful for investigations of matrotrophy, and was used where female-specific data were absent. However, as our clade-wide analyses span multiple orders of magnitude and residuals are generated as log-log interactions, we suggest sex-specific nuances will have little influence on the overall outcome of our work.

In conclusion, by demonstrating the interaction of matrotrophic investment and reduced longevity across large vertebrate clades, we have moved the understanding of DST dynamics beyond an explanation for individual senescence to one of a key trade-off defining the evolution of reproductive life history strategies across species. We find support for the DST uniquely associated with the origin and degree of viviparous matrotrophy at a macroevolutionary scale, with no evidence that the SMH acts as an evolutionary driver. As such, we propose this specific framework of reproductive diversification across the vertebrates as the Sacrificial Mother model, where longevity is reduced by the sum of maternal costs from high matrotrophy traded off against increased embryonic fitness. By investigating this phenomenon in both chondrichthyans and mammals, controlled for both body size and phylogeny, we show that Sacrificial Mothers are a signature of both convergent and divergent reproductive evolution and associated with both the qualitative origin and quantitative degree of viviparous matrotrophy. Our united framework decouples parity mode from nutrient provisioning strategy and proposes that the same innovation of matrotrophic investment which elevates offspring fitness decreases maternal lifespan across entire clades. We therefore present Sacrificial Mother dynamics as a fundamental and previously unrecognised driver of vertebrate life history evolution.

## Acknowledgements

We are grateful to Dr. Deby Cassill (University of South Florida) for sharing one of the datasets used to compile chondrichthyan life history metrics in our analysis. RL and DL were funded by BBSRC grant number BB/Y005953/1.

## Supplemental information

All data and code used in this study are freely available as Supplementary Files alongside this manuscript.

