## Supplementary material for "Sacrificial mothers: increased matrotrophy is associated with reduced maternal longevity across chondrichthyans and mammals": Supplementary_File_3_LH_Analysis_Code.R

```
###CHONDRICHTHYAN ANALYSIS###############################################################################

#Load required packages
library(tidyverse)
library(ape)
library(nlme)
library(phytools)
library(car)
library(FSA)
library(patchwork)

#Load shark data
shark_data <- read.csv("### Path to Supplementary_File_1.csv")
shark_tree <- read.nexus("### Path to Shark_Tree.nex")

#Inspect shark data structure
names(shark_data)
str(shark_data)

#Replace any zero/negative branch lengths with a very small positive value
shark_tree$edge.length[shark_tree$edge.length <= 0] <- 1e-6

#Log the shark data and combine reproductive categories 
shark_data <- shark_data %>%
  filter(
    Maximum_longevity_years > 0,
    Maximum_length_cm > 0
  ) %>%
  mutate(
    Log_longevity = log(Maximum_longevity_years),
    Log_length = log(Maximum_length_cm),
    Repro_Strategy = case_when(
      Parity_mode == "Viviparity" & Nutritional_mode == "Lecithotrophy" ~ "Vivip_Lecitho",
      Parity_mode == "Viviparity" & Nutritional_mode == "Matrotrophy" ~ "Vivip_Matro",
      Parity_mode == "Oviparity" & Nutritional_mode == "Lecithotrophy" ~ "Ovip_Lecitho",
      TRUE ~ NA_character_
    )
  )
summary(shark_data[, c("Maximum_longevity_years", "Maximum_length_cm", "Log_longevity", "Log_length")])

#Fit a simple regression to the data
lm1 <- lm(Log_longevity ~ Log_length, data = shark_data)
summary(lm1)
shark_data$resid_longevity <- resid(lm1)

Repro_vec <- setNames(as.character(shark_data$Repro_Strategy), shark_data$Species)

#Quick plot of the simple regression
ggplot(shark_data, aes(x = Log_length, y = Log_longevity)) +
  geom_point() +
  geom_smooth(method = "lm", se = TRUE) +
  theme_bw()

#Inspect the distribution of simple longevity residuals
ggplot(shark_data, aes(x = resid_longevity)) +
  geom_histogram(bins = 30) +
  theme_bw()

#Fit in the loaded phylogenetic tree 
cor_struct <- corPagel(
  value = 0.5,
  phy = shark_tree,
  fixed = FALSE,
  form = ~ Species
)

#Fit baseline PGLS model
pgls_model <- gls(
  Log_longevity ~ Log_length,
  data = shark_data,
  correlation = cor_struct,
  method = "ML"
)

summary(pgls_model)
shark_data$resid_longevity_pgls <- resid(pgls_model)

#Compare the basic and phylogenetic-corrected models
AIC(lm1, pgls_model)
pgls_model$modelStruct$corStruct
coef(pgls_model$modelStruct$corStruct, unconstrained = FALSE)
delta <- AIC(lm1) - AIC(pgls_model)
print(delta)

#Inspect the residual distribution from PGLS model
ggplot(shark_data, aes(x = resid_longevity_pgls)) +
  geom_histogram(bins = 30) +
  theme_bw()

#Create prediction lines for both LM and PGLS for plotting
newdat <- data.frame(
  Log_length = seq(
    min(shark_data$Log_length),
    max(shark_data$Log_length),
    length.out = 200
  )
)

#LM predictions
newdat$pred_lm <- predict(lm1, newdata = newdat)

#PGLS predictions
coef_pgls <- coef(pgls_model)
newdat$pred_pgls <- coef_pgls[1] + coef_pgls[2] * newdat$Log_length

#Plot different models and false colour points by reproductive strategy
repro_cols <- c(
  "Vivip_Lecitho" = "cyan",
  "Vivip_Matro"   = "red",
  "Ovip_Lecitho"  = "white"
)

regression_plot <- ggplot(shark_data, aes(x = Log_length, y = Log_longevity, colour = Repro_Strategy)) +
  geom_point(alpha = 1, size = 3.5) +
  geom_line(
    data = newdat,
    aes(x = Log_length, y = pred_lm),
    inherit.aes = FALSE,
    colour = "black",
    linewidth = 1.1,
    linetype = "dashed"
  ) +
  geom_line(
    data = newdat,
    aes(x = Log_length, y = pred_pgls),
    inherit.aes = FALSE,
    colour = "red",
    linewidth = 1.2
  ) +
  scale_colour_manual(
    values = repro_cols,
    na.value = "grey70"
  ) +
  theme_bw() +
  labs(
    x = "Log maximum length",
    y = "Log maximum longevity",
    colour = "Reproductive strategy"
  )

regression_plot

#Fit the PGLS model and add the residuals
pgls_model <- gls(
  Log_longevity ~ Log_length,
  data = shark_data,
  correlation = corPagel(0.5, phy = shark_tree, fixed = FALSE, form = ~ Species),
  method = "REML"
)
shark_data$resid_longevity_pgls <- resid(pgls_model)

#Test assumptions and differences of life history variables by reproductive strategy
vars <- c(Log_length = "Log maximum length",
          Log_longevity = "Log maximum longevity",
          resid_longevity_pgls = "Relative longevity (PGLS residuals)")

test_var <- function(v){
  f <- reformulate("Repro_Strategy", v)
  shap <- shark_data %>% group_by(Repro_Strategy) %>%
    summarise(shapiro_p = shapiro.test(.data[[v]])$p.value, .groups = "drop")
  lev <- car::leveneTest(f, data = shark_data)$`Pr(>F)`[1]
  param <- all(shap$shapiro_p > .05) & lev > .05
  main <- if(param) summary(aov(f, data = shark_data)) else kruskal.test(f, data = shark_data)
  post <- if(param) TukeyHSD(aov(f, data = shark_data)) else FSA::dunnTest(f, data = shark_data, method = "bonferroni")
  list(variable = v, shapiro = shap, levene_p = lev, parametric = param, main = main, posthoc = post)
}

results <- setNames(lapply(names(vars), test_var), names(vars))
results

#Plot the data
plot_var <- function(v, ylab) ggplot(shark_data, aes(Repro_Strategy, .data[[v]], fill = Repro_Strategy)) +
  geom_boxplot(alpha = .5, outlier.shape = NA, colour = "black") +
  geom_jitter(shape = 21, width = .12, size = 3.5, colour = "black", stroke = 1) +
  scale_fill_manual(values = repro_cols) +
  theme_bw() +
  labs(x = "Reproductive strategy", y = ylab, fill = "Reproductive strategy")

plots <- imap(vars, ~plot_var(.y, .x))
plots$Log_length / plots$Log_longevity / plots$resid_longevity_pgls

#Make the contmap of longevity residuals over the phylogenetic tree
resid_vec <- setNames(shark_data$resid_longevity_pgls, shark_data$Species)
resid_vec <- resid_vec[shark_tree$tip.label]
all(names(resid_vec) == shark_tree$tip.label)
thr <- 1
fan_map <- contMap(shark_tree, resid_vec, plot = FALSE, lims = c(-thr, thr))
fan_map <- setMap(fan_map, colors = colorRampPalette(c("blue", "white", "red"))(100))

#Abbreviate the tip labels for the plot 
abbr_species <- function(x) {
  x2 <- gsub("_", " ", x)
  parts <- strsplit(x2, " ")[[1]]
  if (length(parts) >= 2) {
    paste0(substr(parts[1], 1, 1), ". ", parts[2])
  } else {
    x2
  }
}
fan_map$tree$tip.label <- sapply(fan_map$tree$tip.label, abbr_species)

#Set font 
par(family = "Arial")

#Plot contmap without tip labels
plot(
  fan_map,
  type = "fan",
  fsize = 0.25,
  ftype = "off",
  lwd = 3,
  legend = 50,
  outline = TRUE,
  cols = setNames(fan_map$cols, names(fan_map$cols)),
  x.lim = c(-1.6, 1.6) * max(nodeHeights(fan_map$tree)),
  y.lim = c(-1.6, 1.6) * max(nodeHeights(fan_map$tree))
)

#Get plotted tip coordinates
pp <- get("last_plot.phylo", envir = .PlotPhyloEnv)
tip_x <- pp$xx[1:Ntip(fan_map$tree)]
tip_y <- pp$yy[1:Ntip(fan_map$tree)]

#Tip angles
theta <- atan2(tip_y, tip_x)
theta <- ifelse(theta < 0, theta + 2*pi, theta)

#Ring order
ord <- order(theta)
tip_x <- tip_x[ord]
tip_y <- tip_y[ord]
theta <- theta[ord]
species_ord <- fan_map$tree$tip.label[ord]
Repro_ord <- Repro_vec[shark_tree$tip.label][ord]

#Ring radii
r_tip   <- max(sqrt(tip_x^2 + tip_y^2))
r_inner <- r_tip * 0.99
r_outer <- r_tip * 1.07
r_label <- r_tip * 1.08

#Make contMap tips flush to a perfect circle
tt <- seq(0, 2*pi, length.out = 500)
polygon(
  x = c(par("usr")[1], par("usr")[2], rev(r_tip * cos(tt))),
  y = c(par("usr")[4], par("usr")[4], rev(r_tip * sin(tt))),
  col = "white",
  border = NA
)

#Redraw circular outer edge in black
symbols(
  0, 0, circles = r_tip, inches = FALSE, add = TRUE,
  fg = "black", bg = NA, lwd = 1.2
)

repro_col_vec <- repro_cols[Repro_ord]

#Segment widths
dtheta <- c(diff(theta), theta[1] + 2*pi - theta[length(theta)])
theta_start <- theta - dtheta/2
theta_end   <- theta + dtheta/2

#Draw annulus sector
draw_ring_segment <- function(t1, t2, r1, r2, col, n = 30) {
  tt_outer <- seq(t1, t2, length.out = n)
  tt_inner <- seq(t2, t1, length.out = n)
  x <- c(r2 * cos(tt_outer), r1 * cos(tt_inner))
  y <- c(r2 * sin(tt_outer), r1 * sin(tt_inner))
  polygon(x, y, col = col, border = "black", lwd = 0.5)
}

#Draw reproductive strategy ring
for (i in seq_along(theta)) {
  draw_ring_segment(theta_start[i], theta_end[i], r_inner, r_outer, repro_col_vec[i])
}

#Ring outlines in black
symbols(
  0, 0, circles = r_inner, inches = FALSE, add = TRUE,
  fg = "black", bg = NA, lwd = 1.2
)
symbols(
  0, 0, circles = r_outer, inches = FALSE, add = TRUE,
  fg = "black", bg = NA, lwd = 1.2
)

#Add tip labels outside the ring
for (i in seq_along(theta)) {
  angle_deg <- theta[i] * 180 / pi
  x_lab <- r_label * cos(theta[i])
  y_lab <- r_label * sin(theta[i])
  
  if (angle_deg > 90 && angle_deg < 270) {
    text(
      x_lab, y_lab,
      labels = species_ord[i],
      srt = angle_deg + 180,
      adj = 1,
      cex = 0.25,
      xpd = TRUE,
      font = 3,
      family = "Arial"
    )
  } else {
    text(
      x_lab, y_lab,
      labels = species_ord[i],
      srt = angle_deg,
      adj = 0,
      cex = 0.25,
      xpd = TRUE,
      font = 3,
      family = "Arial"
    )
  }
}

###MAMMAL ANALYSIS###############################################################################

#Load mammal data
mammal_data <- read.csv("### Path to Supplementary_File_2.csv")
mammal_tree <- read.nexus("### Path to Mammal_Tree.nex")

#Inspect the mammal data
names(mammal_data)
str(mammal_data)

#Replace any zero/negative branch lengths with a very small positive value
mammal_tree$edge.length[mammal_tree$edge.length <= 0] <- 1e-6

#Log the mammal data
mammal_data <- mammal_data %>%
  filter(
    Lifespan_days > 0,
    Adult_mass_kg > 0
  ) %>%
  mutate(
    Log_longevity = log(Lifespan_days),
    Log_mass = log(Adult_mass_kg)
  )

#Fit a basic regression and calculate residuals (for exploration only)
lm1 <- lm(Log_longevity ~ Log_mass, data = mammal_data)
summary(lm1)
mammal_data$resid_longevity <- resid(lm1)

#Do a quick plot of the simple regression 
ggplot(mammal_data, aes(x = Log_mass, y = Log_longevity)) +
  geom_point() +
  geom_smooth(method = "lm", se = TRUE) +
  theme_bw()

#Inspect distribution of simple residuals 
ggplot(mammal_data, aes(x = resid_longevity)) +
  geom_histogram(bins = 30) +
  theme_bw()

#Fit in the tree for the phylogenetic model
cor_struct <- corPagel(
  value = 0.5,
  phy = mammal_tree,
  fixed = FALSE,
  form = ~ Species
)

#Fit baseline PGLS model
pgls_model <- gls(
  Log_longevity ~ Log_mass,
  data = mammal_data,
  correlation = cor_struct,
  method = "ML"
)

mammal_data$resid_longevity_pgls <- resid(pgls_model)
summary(pgls_model)

#Compare the simple regression against the phylogenetic model
AIC(lm1, pgls_model)
pgls_model$modelStruct$corStruct
coef(pgls_model$modelStruct$corStruct, unconstrained = FALSE)
delta <- AIC(lm1) - AIC(pgls_model)
print(delta)

#Look at the distribution of residuals run from the PGLS model
ggplot(mammal_data, aes(x = resid_longevity_pgls)) +
  geom_histogram(bins = 30) +
  theme_bw()

#Create prediction lines for both LM and PGLS for plotting
newdat <- data.frame(
  Log_mass = seq(
    min(mammal_data$Log_mass),
    max(mammal_data$Log_mass),
    length.out = 200
  )
)

newdat$pred_lm <- predict(lm1, newdata = newdat)
newdat$pred_pgls <- predict(pgls_model, newdata = newdat)

#Plot different models and colour points by MI
regression_plot <- ggplot(mammal_data, aes(x = Log_mass, y = Log_longevity)) +
  geom_point(aes(colour = MI), alpha = 1, size = 3.5) +
  geom_line(
    data = newdat,
    aes(x = Log_mass, y = pred_lm, linetype = "LM"),
    colour = "black",
    linewidth = 1
  ) +
  geom_line(
    data = newdat,
    aes(x = Log_mass, y = pred_pgls, linetype = "PGLS"),
    colour = "red",
    linewidth = 1
  ) +
  scale_colour_gradientn(
    colours = c("blue", "cyan", "green", "yellow", "orange", "red"),
    name = "MI"
  ) +
  scale_linetype_manual(
    values = c("LM" = "dashed", "PGLS" = "solid"),
    name = "Model"
  ) +
  theme_bw() +
  labs(
    x = "Log adult mass",
    y = "Log lifespan"
  )

regression_plot

#Investigate whether MI is associated with absolute lifespan and body mass

#Lifespan against MI (simple regression)
lm_longevity_MI <- lm(Log_longevity ~ MI, data = mammal_data)
summary(lm_longevity_MI)
#Lifespan against MI (PGLS)
pgls_longevity_MI <- gls(
  Log_longevity ~ MI,
  data = mammal_data,
  correlation = corPagel(0.5, phy = mammal_tree, form = ~ Species),
  method = "ML"
)
summary(pgls_longevity_MI)

#Mass against MI (simple regression)
lm_mass_MI <- lm(Log_mass ~ MI, data = mammal_data)
summary(lm_mass_MI)
#Mass against MI (PGLS)
pgls_mass_MI <- gls(
  Log_mass ~ MI,
  data = mammal_data,
  correlation = corPagel(0.5, phy = mammal_tree, form = ~ Species),
  method = "ML"
)
summary(pgls_mass_MI)

#Quick plots of absolute relationships

#Plot lifespan against MI
ggplot(mammal_data, aes(x = MI, y = Log_longevity)) +
  geom_point(alpha = 1, size = 2.25) +
  geom_smooth(method = "lm", se = TRUE) +
  theme_bw() +
  labs(
    x = "MI",
    y = "Log lifespan",
    title = "Relationship between MI and absolute lifespan"
  )

#Plot mass against MI
ggplot(mammal_data, aes(x = MI, y = Log_mass)) +
  geom_point(alpha = 1, size = 2.25) +
  geom_smooth(method = "lm", se = TRUE) +
  theme_bw() +
  labs(
    x = "MI",
    y = "Log adult mass",
    title = "Relationship between MI and adult body mass"
  )

#Compare the relationship between residual longevity and MI using simple and phylogenetically controlled tests

#Simple residuals vs MI
cor.test(mammal_data$resid_longevity_pgls, mammal_data$MI, method = "pearson")
lm_resid_mi <- lm(resid_longevity_pgls ~ MI, data = mammal_data)

#Phylogenetically controlled model on residuals vs MI
pgls_resid_mi <- gls(
  resid_longevity_pgls ~ MI,
  data = mammal_data,
  correlation = corPagel(0.5, phy = mammal_tree, form = ~ Species),
  method = "ML"
)

summary(lm_resid_mi)
summary(pgls_resid_mi)

#Test phylogenetically controlled correlation in the full longevity model
pgls_full <- gls(
  Log_longevity ~ Log_mass + MI,
  data = mammal_data,
  correlation = corPagel(0.5, phy = mammal_tree, form = ~ Species),
  method = "REML"
)
summary(pgls_full)

#Make prediction grid for plotting the two fitted residual lines
pred_df <- data.frame(
  MI = seq(min(mammal_data$MI), max(mammal_data$MI), length.out = 200)
)

pred_df$lm_fit   <- predict(lm_resid_mi, newdata = pred_df)
pred_df$pgls_fit <- predict(pgls_resid_mi, newdata = pred_df)

#Plot MI against longevity residuals with LM and PGLS lines
mi_residual_plot <- ggplot(mammal_data, aes(x = MI, y = resid_longevity_pgls)) +
  geom_point(
    aes(fill = MI),
    shape = 21,
    size = 3.5,
    color = "black",
    stroke = 1,
    alpha = 1
  ) +
  geom_line(
    data = pred_df,
    aes(x = MI, y = lm_fit),
    color = "black",
    linewidth = 1,
    linetype = "dashed"
  ) +
  geom_line(
    data = pred_df,
    aes(x = MI, y = pgls_fit),
    color = "blue",
    linewidth = 1.1
  ) +
  scale_fill_gradientn(
    colours = c("blue", "cyan", "green", "yellow", "orange", "red")
  ) +
  theme_bw() +
  labs(
    x = "MI",
    y = "Longevity residuals (PGLS)",
    title = "Relationship between MI and longevity residuals",
    fill = "MI"
  )

mi_residual_plot

#Plot contmap to visualise residuals over phylogeny
resid_vec <- setNames(mammal_data$resid_longevity_pgls, mammal_data$Species)
resid_vec <- resid_vec[mammal_tree$tip.label]
resid_vec <- resid_vec[!is.na(resid_vec)]

MI_vec <- setNames(mammal_data$MI, mammal_data$Species)
MI_vec <- MI_vec[names(resid_vec)]

tree2 <- drop.tip(mammal_tree, setdiff(mammal_tree$tip.label, names(resid_vec)))
resid_vec <- resid_vec[tree2$tip.label]
MI_vec <- MI_vec[tree2$tip.label]

#Residual colour threshold 
thr <- 1
fan_map <- contMap(tree2, resid_vec, plot = FALSE, lims = c(-thr, thr))
fan_map <- setMap(fan_map, colors = colorRampPalette(c("blue", "white", "red"))(100))

#Abbreviate tip labels: Genus_species -> G. species 
abbr_species <- function(x) {
  x2 <- gsub("_", " ", x)
  parts <- strsplit(x2, " ")[[1]]
  if (length(parts) >= 2) {
    paste0(substr(parts[1], 1, 1), ". ", parts[2])
  } else {
    x2
  }
}

fan_map$tree$tip.label <- sapply(fan_map$tree$tip.label, abbr_species)

#Plot contmap without tip labels
plot(
  fan_map,
  type = "fan",
  fsize = 0.15,
  ftype = "off",   # <- suppress built-in tip labels
  lwd = 2,
  legend = 50,
  outline = TRUE,
  cols = setNames(fan_map$cols, names(fan_map$cols)),
  x.lim = c(-1.6, 1.6) * max(nodeHeights(tree2)),
  y.lim = c(-1.6, 1.6) * max(nodeHeights(tree2))
)

#Get plotted tip coordinates
pp <- get("last_plot.phylo", envir = .PlotPhyloEnv)
tip_x <- pp$xx[1:Ntip(fan_map$tree)]
tip_y <- pp$yy[1:Ntip(fan_map$tree)]

#Tip angles
theta <- atan2(tip_y, tip_x)
theta <- ifelse(theta < 0, theta + 2 * pi, theta)

#Ring order
ord <- order(theta)
tip_x <- tip_x[ord]
tip_y <- tip_y[ord]
theta <- theta[ord]
species_ord <- fan_map$tree$tip.label[ord]   # <- use abbreviated labels
MI_ord <- MI_vec[tree2$tip.label][ord]       # keep MI in same plotting order

#Radii
r_tip   <- max(sqrt(tip_x^2 + tip_y^2))
r_inner <- r_tip * 0.96      # <- ring starts immediately at contmap edge
r_outer <- r_tip * 1.07      # adjust thickness of ring here
r_label <- r_tip * 1.08      # labels outside the ring

#Make contMap tips flush to a perfect circle
tt <- seq(0, 2 * pi, length.out = 500)
polygon(
  x = c(par("usr")[1], par("usr")[2], rev(r_tip * cos(tt))),
  y = c(par("usr")[4], par("usr")[4], rev(r_tip * sin(tt))),
  col = "white",
  border = NA
)

#Redraw circular outer edge in black
symbols(
  0, 0, circles = r_tip, inches = FALSE, add = TRUE,
  fg = "black", bg = NA, lwd = 1.2
)

#MI colour threshold (adjust min max values)
mi_min <- -0.5
mi_max <- 0.8

#Cap MI values at the chosen threshold
MI_ord_plot <- pmax(pmin(MI_ord, mi_max), mi_min)

#MI colour scale using fixed manual limits
mi_cols <- colorRampPalette(c("navy", "cyan", "yellow", "red"))(100)
mi_breaks <- seq(mi_min, mi_max, length.out = 101)
mi_col_vec <- mi_cols[findInterval(MI_ord_plot, mi_breaks, all.inside = TRUE)]

#Segment widths
dtheta <- c(diff(theta), theta[1] + 2 * pi - theta[length(theta)])
theta_start <- theta - dtheta / 2
theta_end   <- theta + dtheta / 2

#Draw annulus sector
draw_ring_segment <- function(t1, t2, r1, r2, col, n = 30) {
  tt_outer <- seq(t1, t2, length.out = n)
  tt_inner <- seq(t2, t1, length.out = n)
  x <- c(r2 * cos(tt_outer), r1 * cos(tt_inner))
  y <- c(r2 * sin(tt_outer), r1 * sin(tt_inner))
  polygon(x, y, col = col, border = "black", lwd = 0.5)
}

#Draw MI ring
for (i in seq_along(theta)) {
  draw_ring_segment(theta_start[i], theta_end[i], r_inner, r_outer, mi_col_vec[i])
}

#Ring outlines
symbols(
  0, 0, circles = r_inner, inches = FALSE, add = TRUE,
  fg = "black", bg = NA, lwd = 1.2
)
symbols(
  0, 0, circles = r_outer, inches = FALSE, add = TRUE,
  fg = "black", bg = NA, lwd = 1.2
)

#Add tip labels to outside of the ring
for (i in seq_along(theta)) {
  angle_deg <- theta[i] * 180 / pi
  x_lab <- r_label * cos(theta[i])
  y_lab <- r_label * sin(theta[i])
  
  #Flip text on left side so it is readable
  if (angle_deg > 90 && angle_deg < 270) {
    text(
      x_lab, y_lab,
      labels = species_ord[i],
      srt = angle_deg + 180,
      adj = 1,
      cex = 0.12,
      xpd = TRUE,
      font = 3   # italic
    )
  } else {
    text(
      x_lab, y_lab,
      labels = species_ord[i],
      srt = angle_deg,
      adj = 0,
      cex = 0.12,
      xpd = TRUE,
      font = 3   # italic
    )
  }
}
```
