## Supplementary material for "Sacrificial mothers: increased matrotrophy is associated with reduced maternal longevity across chondrichthyans and mammals": Supplementary_File_4_Simulation_Code.ipynb

```
# %% [code]
#### Simple stochastic simulation to model Sacrifical Mother and Disposable Womb dynamics ####

import numpy as np
import matplotlib.pyplot as plt
from matplotlib.patches import Circle, Rectangle
from matplotlib.colors import Normalize, LinearSegmentedColormap
from matplotlib.cm import ScalarMappable

# PARAMETERS: Adjust to explore the interaction of variables

#Fixed random seed for reproducibility 
np.random.seed(42)

# Simulation size and length
N_GENERATIONS = 100
TIMEPOINTS = [0, 25, 50, 75, 100]
POPULATION_SIZE = 100

# Embryo investment (the trait which evolves)
INITIAL_INVESTMENT = 0.35

# Investment ceilings
LECITHO_CAP = 1.10
MATROTROPHY_CAP = 1.75

# Matrotrophy-specific damage onset
DAMAGE_THRESHOLD = 1.2

# Mutation rate
INVESTMENT_INCREASE_MEAN = 0.015
MUTATION_SD = 0.02

# Maternal investment effects
MATERNAL_NUTRIENTS = 0.020
POST_CAP_DECAY = 0.06
POST_CAP_PENALTY = 0.040

# Slight boost in investment for viviparous lecithotrophy (maternal control of gestation) 
VIVIP_LECITHO_BONUS = 0.25

# Fitness parameters
MINIMUM_FITNESS = 1e-6
BASE_BENEFIT_SCALE = 1.20
BASE_BENEFIT_STEEPNESS = 2.40

# Additional matrotrophic offspring benefit once exceeding lecithotrophic cap
MATRO_EXTRA_BENEFIT_SCALE = 2.10
MATRO_EXTRA_BENEFIT_STEEPNESS = 4.20

# Protection / retention benefit from internalised gestation 
PROTECTION_BONUS = {
    "Oviparity": 0.00,
    "Viviparous Lecithotrophy": 0.25,
    "Viviparous Matrotrophy": 0.25,
}

# Baseline lifespans different by parity mode (viviparous have higher baseline from larger-bodied slower LHS)
BASELINE_LIFESPAN_BY_STRATEGY = {
    "Oviparity": 12.0,
    "Viviparous Lecithotrophy": 14.0,
    "Viviparous Matrotrophy": 14.0,
}

MINIMUM_MATERNAL_LIFESPAN = 0.2

# Maternal damage
DAMAGE_EXPONENT = 1.5
DAMAGE_SCALE = 4
FITNESS_DAMAGE_COST = 0.30

# Lifespan decline from damage (highly non-linear at extreme values)
LIFESPAN_DAMAGE_SCALE = 0.55
LIFESPAN_DAMAGE_EXPONENT = 1.45
LIFESPAN_EXTREME_THRESHOLD = 1.2
LIFESPAN_EXTREME_RATE = 0.90

# Visualise single-lineage trajectory (size of embryo represent investment, not physical size) 
MOTHER_RADIUS = 3.0
WOMB_RADIUS = 2.1
VISUAL_SCALE = 1.35


### DEFINE STRATEGIES 
strategies = {
    "Oviparity": {
        "investment_max": LECITHO_CAP,
        "nutrient_supply": 0.0,
        "retention_bonus": 0.00,
        "protection_bonus": PROTECTION_BONUS["Oviparity"],
        "damage_enabled": False,
        "baseline_lifespan": BASELINE_LIFESPAN_BY_STRATEGY["Oviparity"],
        "color": "blue",
    },
    "Viviparous Lecithotrophy": {
        "investment_max": LECITHO_CAP + VIVIP_LECITHO_BONUS,
        "nutrient_supply": 0.0,
        "retention_bonus": 0.01,
        "protection_bonus": PROTECTION_BONUS["Viviparous Lecithotrophy"],
        "damage_enabled": False,
        "baseline_lifespan": BASELINE_LIFESPAN_BY_STRATEGY["Viviparous Lecithotrophy"],
        "color": "orange",
    },
    "Viviparous Matrotrophy": {
        "investment_max": MATROTROPHY_CAP,
        "nutrient_supply": MATERNAL_NUTRIENTS,
        "retention_bonus": 0.01,
        "protection_bonus": PROTECTION_BONUS["Viviparous Matrotrophy"],
        "damage_enabled": True,
        "baseline_lifespan": BASELINE_LIFESPAN_BY_STRATEGY["Viviparous Matrotrophy"],
        "color": "green",
    },
}

STRATEGY_ORDER = list(strategies.keys())
METRIC_NAMES = ("investments", "damages", "fitnesses", "lifespans")

### CORE MODEL FUNCTIONS 

def maternal_damage(investment, damage_enabled):
    """
    Maternal damage occurs only in matrotrophy, once investment exceeds
    the safe threshold. Damage rises nonlinearly.
    """
    if not damage_enabled:
        return 0.0

    overshoot = max(0.0, investment - DAMAGE_THRESHOLD)
    return DAMAGE_SCALE * (overshoot ** DAMAGE_EXPONENT)


def maternal_lifespan_from_damage(damage, baseline_lifespan):
    """
    Maternal lifespan declines nonlinearly with damage.
    Small damage has a modest effect; larger damage causes a sharper collapse.
    """
    mild_component = LIFESPAN_DAMAGE_SCALE * (damage ** LIFESPAN_DAMAGE_EXPONENT)
    lifespan = baseline_lifespan - mild_component

    if damage > LIFESPAN_EXTREME_THRESHOLD:
        lifespan *= np.exp(-LIFESPAN_EXTREME_RATE * (damage - LIFESPAN_EXTREME_THRESHOLD))

    return max(MINIMUM_MATERNAL_LIFESPAN, lifespan)


def embryo_fitness(investment, strategy_name, protection_bonus, damage):
    """
    Fitness has:
    - a general saturating benefit of embryo investment
    - a strategy-specific protection bonus
    - an extra steep benefit for matrotrophy above the lecithotrophic ceiling
    - a maternal-damage-associated cost
    """
    base_benefit = BASE_BENEFIT_SCALE * (1.0 - np.exp(-BASE_BENEFIT_STEEPNESS * investment))

    extra_matro_benefit = 0.0
    if strategy_name == "Viviparous Matrotrophy" and investment > LECITHO_CAP:
        excess = investment - LECITHO_CAP
        extra_matro_benefit = MATRO_EXTRA_BENEFIT_SCALE * (
            1.0 - np.exp(-MATRO_EXTRA_BENEFIT_STEEPNESS * excess)
        )

    damage_cost = FITNESS_DAMAGE_COST * damage

    fitness = 1.0 + base_benefit + protection_bonus + extra_matro_benefit - damage_cost
    return max(MINIMUM_FITNESS, fitness)


def propose_investment(current_investment, investment_max, nutrient_supply, retention_bonus):
    """
    Propose a new embryo investment value after mutation and provisioning constraints.

    Oviparity:
        capped at the lecithotrophic ceiling.

    Viviparous lecithotrophy:
        slightly exceeds oviparity because retained development/protection
        incurs a modest added investment cost, but it does not enter the
        strongly damaging matrotrophic regime.

    Viviparous matrotrophy:
        can continue beyond the lecithotrophic ceiling through maternal provisioning.
    """
    change = np.random.normal(INVESTMENT_INCREASE_MEAN, MUTATION_SD) + retention_bonus

    if current_investment >= LECITHO_CAP:
        overshoot = current_investment - LECITHO_CAP
        if nutrient_supply > 0:
            change += nutrient_supply
            change -= POST_CAP_DECAY * overshoot
        else:
            change -= POST_CAP_PENALTY
            change -= POST_CAP_DECAY * overshoot

    candidate = max(0.05, current_investment + change)
    return min(candidate, investment_max)


def accept_candidate(current_investment, candidate_investment, strategy_name, protection_bonus, damage_enabled):
    """
    Accept fitter candidates always; accept worse ones probabilistically.
    """
    current_damage = maternal_damage(current_investment, damage_enabled)
    candidate_damage = maternal_damage(candidate_investment, damage_enabled)

    current_fitness = embryo_fitness(current_investment, strategy_name, protection_bonus, current_damage)
    candidate_fitness = embryo_fitness(candidate_investment, strategy_name, protection_bonus, candidate_damage)

    if candidate_fitness >= current_fitness:
        return candidate_investment, candidate_damage, candidate_fitness

    acceptance_prob = candidate_fitness / current_fitness
    if np.random.rand() < acceptance_prob:
        return candidate_investment, candidate_damage, candidate_fitness

    return current_investment, current_damage, current_fitness


def simulate_embryo_investment(
    n_generations,
    initial_investment,
    investment_max,
    strategy_name,
    nutrient_supply=0.0,
    retention_bonus=0.0,
    protection_bonus=0.0,
    damage_enabled=False,
    baseline_lifespan=12.0,
):
    """Simulate one embryo-investment trajectory across generations."""
    investments = np.zeros(n_generations + 1)
    damages = np.zeros(n_generations + 1)
    fitnesses = np.zeros(n_generations + 1)
    lifespans = np.zeros(n_generations + 1)

    investments[0] = initial_investment
    damages[0] = maternal_damage(initial_investment, damage_enabled)
    fitnesses[0] = embryo_fitness(initial_investment, strategy_name, protection_bonus, damages[0])
    lifespans[0] = maternal_lifespan_from_damage(damages[0], baseline_lifespan)

    for t in range(1, n_generations + 1):
        current_investment = investments[t - 1]
        candidate_investment = propose_investment(
            current_investment=current_investment,
            investment_max=investment_max,
            nutrient_supply=nutrient_supply,
            retention_bonus=retention_bonus,
        )

        new_investment, new_damage, new_fitness = accept_candidate(
            current_investment=current_investment,
            candidate_investment=candidate_investment,
            strategy_name=strategy_name,
            protection_bonus=protection_bonus,
            damage_enabled=damage_enabled,
        )

        investments[t] = new_investment
        damages[t] = new_damage
        fitnesses[t] = new_fitness
        lifespans[t] = maternal_lifespan_from_damage(new_damage, baseline_lifespan)

    return {
        "investments": investments,
        "damages": damages,
        "fitnesses": fitnesses,
        "lifespans": lifespans,
    }


def simulate_population(
    population_size,
    n_generations,
    initial_investment,
    investment_max,
    strategy_name,
    nutrient_supply=0.0,
    retention_bonus=0.0,
    protection_bonus=0.0,
    damage_enabled=False,
    baseline_lifespan=12.0,
):
    """Run many independent embryo-investment trajectories."""
    population = {
        metric: np.zeros((population_size, n_generations + 1))
        for metric in METRIC_NAMES
    }

    for i in range(population_size):
        run = simulate_embryo_investment(
            n_generations=n_generations,
            initial_investment=initial_investment,
            investment_max=investment_max,
            strategy_name=strategy_name,
            nutrient_supply=nutrient_supply,
            retention_bonus=retention_bonus,
            protection_bonus=protection_bonus,
            damage_enabled=damage_enabled,
            baseline_lifespan=baseline_lifespan,
        )
        for metric in METRIC_NAMES:
            population[metric][i] = run[metric]

    return population


def summarize_population(matrix):
    """Return mean and central 95% interval across trajectories."""
    return {
        "mean": np.mean(matrix, axis=0),
        "low": np.percentile(matrix, 2.5, axis=0),
        "high": np.percentile(matrix, 97.5, axis=0),
    }


def build_population_summary(population_results):
    """Summarize all population runs for all strategies."""
    summary = {}
    for name, res in population_results.items():
        summary[name] = {metric: summarize_population(res[metric]) for metric in METRIC_NAMES}
        summary[name]["color"] = res["color"]
    return summary


### RUN SINGLE SIMULATIONS
results = {}
for name, params in strategies.items():
    results[name] = simulate_embryo_investment(
        n_generations=N_GENERATIONS,
        initial_investment=INITIAL_INVESTMENT,
        investment_max=params["investment_max"],
        strategy_name=name,
        nutrient_supply=params["nutrient_supply"],
        retention_bonus=params["retention_bonus"],
        protection_bonus=params["protection_bonus"],
        damage_enabled=params["damage_enabled"],
        baseline_lifespan=params["baseline_lifespan"],
    )

generations = np.arange(N_GENERATIONS + 1)

### PLOT HELPERS
def plot_metric_lines(ax, x, results_dict, metric, ylabel, title):
    """Plot a single metric across generations for all strategies."""
    for name in STRATEGY_ORDER:
        ax.plot(
            x,
            results_dict[name][metric],
            color=strategies[name]["color"],
            linewidth=2.2,
            label=f"{name} {ylabel.lower()}",
        )
    ax.set_xlabel("Generation")
    ax.set_ylabel(ylabel)
    ax.set_title(title)
    ax.grid(True, alpha=0.3)
    ax.legend()


def plot_population_metric(ax, x, population_summary, metric, ylabel, title):
    """Plot mean trajectory and 95% interval for each strategy."""
    for name in STRATEGY_ORDER:
        summary = population_summary[name]
        color = summary["color"]

        ax.fill_between(
            x,
            summary[metric]["low"],
            summary[metric]["high"],
            color=color,
            alpha=0.18,
        )
        ax.plot(
            x,
            summary[metric]["mean"],
            color=color,
            linewidth=2.5,
            label=f"{name} mean {ylabel.lower()}",
        )

    ax.set_xlabel("Generation")
    ax.set_ylabel(ylabel)
    ax.set_title(title)
    ax.grid(True, alpha=0.3)
    ax.legend()


# Plot investment + damage
fig, ax1 = plt.subplots(figsize=(11, 5))

line_handles = []
damage_handles = []

for name in STRATEGY_ORDER:
    line, = ax1.plot(
        generations,
        results[name]["investments"],
        color=strategies[name]["color"],
        linewidth=2.5,
        label=f"{name} embryo investment",
    )
    line_handles.append(line)

limit_handles = [
    ax1.axhline(LECITHO_CAP, linestyle="-", linewidth=1.8, color="brown", label="Lecithotrophic ceiling"),
    ax1.axhline(DAMAGE_THRESHOLD, linestyle="-.", linewidth=1.5, color="purple", label="Damage threshold"),
    ax1.axhline(MATROTROPHY_CAP, linestyle="--", linewidth=1.5, color="darkgreen", label="Matrotrophic upper ceiling"),
]

ax1.set_xlabel("Generation")
ax1.set_ylabel("Embryo investment")

ax2 = ax1.twinx()
damage_styles = {
    "Oviparity": "--",
    "Viviparous Lecithotrophy": "-.",
    "Viviparous Matrotrophy": "-",
}

for name in STRATEGY_ORDER:
    dline, = ax2.plot(
        generations,
        results[name]["damages"],
        color="red",
        linewidth=2,
        linestyle=damage_styles[name],
        label=f"{name} maternal damage",
    )
    damage_handles.append(dline)

ax2.set_ylabel("Maternal damage")

handles = line_handles + limit_handles + damage_handles
labels = [h.get_label() for h in handles]
ax1.legend(handles, labels, loc="upper left", fontsize=8)

ax1.grid(True, alpha=0.3)
ax1.set_title("Embryo investment evolution and maternal damage")
plt.tight_layout()
plt.show()

# Plot fitness and lifespan
fig, ax = plt.subplots(figsize=(11, 4))
plot_metric_lines(
    ax,
    generations,
    results,
    metric="fitnesses",
    ylabel="Fitness",
    title="Fitness across generations",
)
plt.tight_layout()
plt.show()

fig, ax = plt.subplots(figsize=(11, 4))
plot_metric_lines(
    ax,
    generations,
    results,
    metric="lifespans",
    ylabel="Maternal lifespan",
    title="Maternal lifespan across generations",
)
plt.tight_layout()
plt.show()

# Damage colormap
damage_cmap = LinearSegmentedColormap.from_list(
    "damage_map",
    [(1, 1, 1), (0, 0, 1), (0, 1, 0), (1, 1, 0), (1, 0, 0)],
    N=256,
)

all_damage = np.concatenate([results[name]["damages"] for name in STRATEGY_ORDER])
damage_norm = Normalize(vmin=np.min(all_damage), vmax=max(np.max(all_damage), 1.0))


def radial_gradient(size=300):
    """Create a circular radial gradient mask."""
    x = np.linspace(-1, 1, size)
    y = np.linspace(-1, 1, size)
    X, Y = np.meshgrid(x, y)
    R = np.sqrt(X**2 + Y**2)
    return np.clip(1 - R, 0, 1)


GRADIENT_MASK = radial_gradient()

# Schematic drawing
def draw_scenario(ax, investment, damage, lifespan, fitness, mode, t):
    """
    Draw a conceptual cross-section.
    The embryo circle represents investment level conceptually, not literal size.
    """
    ax.set_aspect("equal")
    ax.axis("off")

    embryo_color = "#a5d6a7"
    womb_fill = "#f8bbd0"

    halo_color = damage_cmap(damage_norm(damage))
    rgba = np.zeros((GRADIENT_MASK.shape[0], GRADIENT_MASK.shape[1], 4))
    rgba[..., :3] = halo_color[:3]
    rgba[..., 3] = 0.5 * GRADIENT_MASK

    extent = [-MOTHER_RADIUS, MOTHER_RADIUS, -MOTHER_RADIUS, MOTHER_RADIUS]
    im = ax.imshow(rgba, extent=extent, origin="lower")
    im.set_clip_path(Circle((0, 0), MOTHER_RADIUS, transform=ax.transData))

    ax.add_patch(Circle((0, 0), WOMB_RADIUS, facecolor=womb_fill, edgecolor="black", linewidth=2))
    ax.add_patch(Circle((0, 0), MOTHER_RADIUS, facecolor="none", edgecolor="black", linewidth=2))

    embryo_radius = 0.55 + (investment / MATROTROPHY_CAP) * 1.2

    if mode == "Oviparity":
        ax.add_patch(
            Circle(
                (0, 0),
                embryo_radius + 0.12,
                facecolor="none",
                edgecolor="black",
                linewidth=2,
            )
        )

    ax.add_patch(Circle((0, 0), embryo_radius, facecolor=embryo_color, edgecolor="black", linewidth=2))

    # Investment gauge
    bar_x0, bar_y0 = -2.1, -MOTHER_RADIUS - 0.65
    bar_w, bar_h = 4.2, 0.28
    ax.add_patch(Rectangle((bar_x0, bar_y0), bar_w, bar_h, facecolor="white", edgecolor="black", linewidth=1.2))
    fill_w = bar_w * min(investment / MATROTROPHY_CAP, 1.0)
    ax.add_patch(Rectangle((bar_x0, bar_y0), fill_w, bar_h, facecolor="#66bb6a", edgecolor="none"))

    lecitho_x = bar_x0 + bar_w * (LECITHO_CAP / MATROTROPHY_CAP)
    damage_x = bar_x0 + bar_w * (DAMAGE_THRESHOLD / MATROTROPHY_CAP)
    ax.plot([lecitho_x, lecitho_x], [bar_y0 - 0.06, bar_y0 + bar_h + 0.06], color="brown", linewidth=2)
    ax.plot([damage_x, damage_x], [bar_y0 - 0.06, bar_y0 + bar_h + 0.06], color="purple", linewidth=2)

    ax.text(0, bar_y0 - 0.18, "Embryo investment", ha="center", va="top", fontsize=8)
    ax.text(
        0,
        WOMB_RADIUS + 0.15,
        f"Lecithotrophic ceiling = {LECITHO_CAP:.2f}\nDamage threshold = {DAMAGE_THRESHOLD:.2f}",
        ha="center",
        va="bottom",
        fontsize=8,
    )
    ax.text(0, MOTHER_RADIUS + 0.45, f"{mode}\nT = {t}", ha="center", va="bottom", fontsize=10)
    ax.text(
        0,
        -MOTHER_RADIUS - 1.45,
        f"Embryo investment = {investment:.2f}\n"
        f"Maternal damage = {damage:.2f}\n"
        f"Maternal lifespan = {lifespan:.2f}\n"
        f"Fitness = {fitness:.2f}",
        ha="center",
        va="top",
        fontsize=9,
    )

    lim = MOTHER_RADIUS + 1
    ax.set_xlim(-lim, lim)
    ax.set_ylim(-lim - 1.3, lim)

# Plot snapshots
fig, axes = plt.subplots(3, len(TIMEPOINTS), figsize=(3 * len(TIMEPOINTS), 10))

for col, t in enumerate(TIMEPOINTS):
    for row, name in enumerate(STRATEGY_ORDER):
        draw_scenario(
            axes[row, col],
            results[name]["investments"][t],
            results[name]["damages"][t],
            results[name]["lifespans"][t],
            results[name]["fitnesses"][t],
            name,
            t,
        )

fig.suptitle("Schematic snapshots across generations", fontsize=14, y=0.98)
plt.tight_layout()
plt.show()

# Colorbar
fig, ax = plt.subplots(figsize=(6, 1))
fig.subplots_adjust(bottom=0.5)

sm = ScalarMappable(norm=damage_norm, cmap=damage_cmap)
sm.set_array([])

cbar = plt.colorbar(sm, cax=ax, orientation="horizontal")
cbar.set_label("Maternal damage (white → red)")
plt.show()

### RUN POPULATION SIMULATIONS
population_results = {}
for name, params in strategies.items():
    population_results[name] = simulate_population(
        population_size=POPULATION_SIZE,
        n_generations=N_GENERATIONS,
        initial_investment=INITIAL_INVESTMENT,
        investment_max=params["investment_max"],
        strategy_name=name,
        nutrient_supply=params["nutrient_supply"],
        retention_bonus=params["retention_bonus"],
        protection_bonus=params["protection_bonus"],
        damage_enabled=params["damage_enabled"],
        baseline_lifespan=params["baseline_lifespan"],
    )
    population_results[name]["color"] = params["color"]

population_summary = build_population_summary(population_results)

# Population plots
fig, ax = plt.subplots(figsize=(11, 5))
plot_population_metric(
    ax,
    generations,
    population_summary,
    metric="investments",
    ylabel="Embryo investment",
    title="Population embryo investment trajectories",
)
ax.axhline(LECITHO_CAP, color="brown", linestyle="-", linewidth=1.3, alpha=0.8, label="Lecithotrophic ceiling")
ax.axhline(DAMAGE_THRESHOLD, color="purple", linestyle="-.", linewidth=1.3, alpha=0.8, label="Damage threshold")
ax.legend()
plt.tight_layout()
plt.show()

fig, ax = plt.subplots(figsize=(11, 4))
plot_population_metric(
    ax,
    generations,
    population_summary,
    metric="damages",
    ylabel="Maternal damage",
    title="Population maternal damage",
)
plt.tight_layout()
plt.show()

fig, ax = plt.subplots(figsize=(11, 4))
plot_population_metric(
    ax,
    generations,
    population_summary,
    metric="fitnesses",
    ylabel="Fitness",
    title="Population embryo fitness",
)
plt.tight_layout()
plt.show()

fig, ax = plt.subplots(figsize=(11, 4))
plot_population_metric(
    ax,
    generations,
    population_summary,
    metric="lifespans",
    ylabel="Maternal lifespan",
    title="Population maternal lifespan",
)
plt.tight_layout()
plt.show()

# Plot final investment histogram
fig, ax = plt.subplots(figsize=(11, 5))

final_investments = [population_results[name]["investments"][:, -1] for name in STRATEGY_ORDER]
bins = np.linspace(
    min(arr.min() for arr in final_investments),
    max(arr.max() for arr in final_investments),
    18,
)

ax.hist(
    final_investments,
    bins=bins,
    stacked=True,
    color=[strategies[name]["color"] for name in STRATEGY_ORDER],
    edgecolor="black",
    label=["Oviparity", "Lecithotrophy", "Matrotrophy"],
    alpha=0.85,
)

ax.axvline(LECITHO_CAP, color="brown", linestyle="-", linewidth=1.4, alpha=0.8, label="Lecithotrophic ceiling")
ax.axvline(DAMAGE_THRESHOLD, color="purple", linestyle="-.", linewidth=1.4, alpha=0.8, label="Damage threshold")
ax.set_xlabel("Final embryo investment")
ax.set_ylabel("Count")
ax.set_title("Stacked histogram of final embryo investments")
ax.grid(True, alpha=0.25)
ax.legend()

plt.tight_layout()
plt.show()

# Print mean population outputs
print("\n=== Population mean outputs by strategy ===")

for name in STRATEGY_ORDER:
    summary = population_summary[name]
    print(f"\n{name}")
    print("-" * len(name))
    print(f"Final mean embryo investment : {summary['investments']['mean'][-1]:.4f}")
    print(f"Final mean maternal damage   : {summary['damages']['mean'][-1]:.4f}")
    print(f"Final mean embryo fitness    : {summary['fitnesses']['mean'][-1]:.4f}")
    print(f"Final mean maternal lifespan : {summary['lifespans']['mean'][-1]:.4f}")

# %% [code]
#Plot final output figures 
import matplotlib.pyplot as plt
import numpy as np

# Safety check
required_keys = ["investments", "damages", "fitnesses", "lifespans", "color"]
for name, summary in population_summary.items():
    for key in required_keys:
        if key not in summary:
            raise ValueError(f"population_summary['{name}'] is missing key: {key}")

generations = np.arange(N_GENERATIONS + 1)

fig = plt.figure(figsize=(12, 8))
gs = fig.add_gridspec(
    2, 2,
    width_ratios=[1.35, 1.0],
    height_ratios=[1.0, 1.0],
    hspace=0.32,
    wspace=0.28
)

# Panel A: embryo investment through time
axA = fig.add_subplot(gs[:, 0])

for name, summary in population_summary.items():
    color = summary["color"]

    axA.fill_between(
        generations,
        summary["investments"]["low"],
        summary["investments"]["high"],
        color=color,
        alpha=0.16
    )
    axA.plot(
        generations,
        summary["investments"]["mean"],
        color=color,
        linewidth=2.8,
        label=name
    )

axA.axhline(
    LECITHO_CAP,
    color="saddlebrown",
    linestyle="-",
    linewidth=1.4,
    alpha=0.9,
    label="Lecithotrophic ceiling"
)
axA.axhline(
    DAMAGE_THRESHOLD,
    color="purple",
    linestyle="-.",
    linewidth=1.4,
    alpha=0.9,
    label="Damage threshold"
)

axA.set_xlabel("Generation")
axA.set_ylabel("Embryo investment")
axA.set_title("A. Evolution of embryo investment", loc="left", fontweight="bold")
axA.grid(True, alpha=0.25)
axA.legend(frameon=False, fontsize=9)

# Panel B: maternal damage vs fitness
axB = fig.add_subplot(gs[0, 1])

for name, summary in population_summary.items():
    color = summary["color"]

    x_mean = summary["fitnesses"]["mean"]
    y_mean = summary["damages"]["mean"]

    x_low = summary["fitnesses"]["low"]
    x_high = summary["fitnesses"]["high"]
    y_low = summary["damages"]["low"]
    y_high = summary["damages"]["high"]

    # Build a ribbon around the trajectory using paired low/high envelopes
    x_poly = np.concatenate([x_low, x_high[::-1]])
    y_poly = np.concatenate([y_low, y_high[::-1]])

    axB.fill(
        x_poly,
        y_poly,
        color=color,
        alpha=0.14,
        linewidth=0
    )
    axB.plot(
        x_mean,
        y_mean,
        color=color,
        linewidth=2.6,
        label=name
    )
    axB.scatter(
        x_mean[-1],
        y_mean[-1],
        color=color,
        s=42,
        zorder=3
    )

axB.set_xlabel("Embryo fitness")
axB.set_ylabel("Maternal damage")
axB.set_title("B. Maternal cost of embryo fitness", loc="left", fontweight="bold")
axB.grid(True, alpha=0.25)

# Panel C: final evolutionary endpoints
axC = fig.add_subplot(gs[1, 1])

for name, summary in population_summary.items():
    color = summary["color"]

    final_invest_mean = summary["investments"]["mean"][-1]
    final_invest_low = summary["investments"]["low"][-1]
    final_invest_high = summary["investments"]["high"][-1]

    final_life_mean = summary["lifespans"]["mean"][-1]
    final_life_low = summary["lifespans"]["low"][-1]
    final_life_high = summary["lifespans"]["high"][-1]

    xerr = np.array([[final_invest_mean - final_invest_low], [final_invest_high - final_invest_mean]])
    yerr = np.array([[final_life_mean - final_life_low], [final_life_high - final_life_mean]])

    axC.errorbar(
        final_invest_mean,
        final_life_mean,
        xerr=xerr,
        yerr=yerr,
        fmt="o",
        color=color,
        elinewidth=1.6,
        capsize=3,
        markersize=7
    )

    axC.text(
        final_invest_mean + 0.015,
        final_life_mean,
        name,
        color=color,
        fontsize=9,
        va="center"
    )

axC.set_xlabel("Final embryo investment")
axC.set_ylabel("Final maternal lifespan")
axC.set_title("C. Evolutionary endpoints", loc="left", fontweight="bold")
axC.grid(True, alpha=0.25)

# Shared styling
for ax in [axA, axB, axC]:
    ax.spines["top"].set_visible(False)
    ax.spines["right"].set_visible(False)

fig.suptitle(
    "Embryo investment evolution, fitness-associated maternal costs, and final outcomes across reproductive strategies",
    fontsize=14,
    y=0.98
)

plt.tight_layout()

plt.show()
```
